# Balancing tumour proliferation and sustained cell cycle arrest through proteostasis remodelling drives immune niche compartmentalisation in breast cancer

**DOI:** 10.1101/2025.01.08.632014

**Authors:** Cenk Celik, Eloise Withnell, Shi Pan, Tooki Chu, John Labbadia, Maria Secrier

## Abstract

Uncontrolled proliferation is a hallmark of cancer, yet tumour cells can enter G0 arrest by halting the cell cycle reversibly (quiescence) or irreversibly (senescence) to survive under stress and in hostile tumour microenvironments (TME). G0 arrested cells contribute to drug tolerance, metastasis and recurrence, but their identification remains challenging due to their rarity and elusive regulatory pathways. Here, we quantify G0 arrest and proliferation decisions in single-cell and spatially profiled breast primary tumours to unveil adaptive responses driving immune compartmentalisation. We identify a genomically-constrained and prolonged G0 arrest state resembling that of dormant precursors of cancer progression. This state featured adaptive transcriptional reprogramming, including unfolded protein response stress, reduced mitochondrial translation and epithelial-mesenchymal plasticity enabled by semaphorin signalling. Spatial transcriptomics revealed G0 arrest pockets encapsulated by *APOE*+ lipid-associated macrophages, myofibroblastic CAFs and immature perivascular cells, suggestive of an immunosuppressive niche contrasting with cytotoxic environments in proliferative hotspots and displaying distinct drug sensitivities. To facilitate future research, we provide a foundation model capturing G0 arrest and proliferation with 94% accuracy in single cell data, available at https://github.com/secrierlab/G0-LM. Our findings provide new insights into the spatial organisation of cell cycle arrest in breast cancer, highlighting the role of G0 states in tumour heterogeneity and adaptation.

## Introduction

The unchecked proliferation of tumour cells is a hallmark of cancer^1^, driving tumour progression and treatment resistance^2,3^. However, proliferation can halt when cells exit the cell cycle in G0, either reversibly (quiescence or dormancy)^4–6^ or irreversibly (senescence)^7^, often in response to intrinsic or environmental stresses. These pauses allow cancer cells to withstand hostile conditions within the tumour microenvironment (TME), thereby promoting drug tolerance and therapeutic persistence^8,9^. Quiescence, in particular, may either be an inherent trait of drug-tolerant persister cells or an acquired adaptive state^10,11^. During tumour evolution, quiescent cells facilitate immune evasion^12,13^ and niche adaptation^14,15^, fostering minimal residual disease and increasing the risk of relapse^11,16,17^.

Maintaining a balance between proliferation and cell cycle arrest is essential for tumour adaptation. Breast cancer is the quintessential example where cell cycle dynamics underpin clinical outcomes, with receptor-positive subtypes exhibiting slower proliferative capacities, but late recurrence through clinical dormancy, while triple-negative breast cancer (TNBC) is marked by aggressive, rapidly proliferating cells and consequently bears the worst prognosis^18,19^. Epigenetic mechanisms have been implicated in controlling cell cycle switches in ER+ tumours: epithelial-mesenchymal plasticity helps maintain slow proliferation of disseminated tumour cells and prevents reawakening from dormancy^20,21^, while chromatin remodelling aids dormancy-driven resistance to endocrine therapies^22^. Cell cycle activity in breast tumours is also influenced by interactions with the TME, which further impacts therapy outcomes. For instance, Baldominos et al.^23^ used spatial transcriptomics to describe a quiescent, fibroblast-enriched niche capable of evading immune detection in TNBC.

The clinical relevance of G0 arrest extends beyond breast cancer, with reports of slow cycling, revival stem cell phenotypes linked with poor prognosis and chemoresistance in colorectal cancer^24–26^, or quiescent *PROM1*+ cells implicated in recurrence and resistance in paediatric high-grade gliomas^27^. However, studying G0 arrested cells remains challenging due to their diverse phenotypes^28,29^, scarcity within tumours and plastic switching between proliferative and quiescent states. The often short-lived nature of this state complicates its identification, while the signalling pathways controlling G0 arrest are still poorly understood. Additionally, the interplay between intrinsic factors^30,31^ and the tumour ecosystem in shaping G0 arrest and proliferation decisions is crucial in determining invasiveness^32^ and therapy resistance^33–35^, but remains inadequately explored.

We previously developed a robust method to quantify G0 arrest in tumours using bulk or single-cell transcriptomic data, identifying genetic modulators like CEP89 that influence G0 arrest capacity^36^. Here, we apply this method to examine G0 arrest and proliferation phenotypes across breast cancer cohorts at single-cell and spatial transcriptomics resolution. We identify a stress-induced, G0 arrested, hybrid epithelial-mesenchymal (E/M) state in primary breast tumours, which appears longer-lived, presents unfolded protein response (UPR) and is linked to immune evasion via the semaphorin/plexin axis. We further uncover spatially segregated G0 arrest and proliferative niches throughout tumours with distinct TME dynamics and drug efficacy. Finally, we develop a foundation model to facilitate the identification of G0, slow- and fast-cycling states in single cell RNA-seq breast cancer data, available at https://github.com/secrierlab/G0-LM. Our findings emphasise the pervasive role of G0 arrest in breast cancer, providing a foundation for future research into its implications to tumour adaptation, progression and the cellular diversity that shapes breast cancer phenotypes.

## Results

### The landscape of G0 arrest and proliferation decisions in primary breast tumours

We collated 138,727 cells across various breast cancer subtypes, leveraging 10X Genomics single-cell RNA sequencing (scRNA-seq) datasets from three studies^37–39^ deposited at the Curated Cancer Cell Atlas of the Weizmann Institute of Science^40^. Our atlas covered both non-invasive (including ductal carcinoma *in situ* [DCIS] and neoplastic) and invasive breast cancers— hormone-positive (ER+ or PR+), HER2+ and triple-negative breast cancer (TNBC)—from 43 patients, and four healthy individuals (Fig. 1a-c; Extended Data Fig. 1a-c). A heterogeneous landscape of cell populations was observed across these tumours and healthy tissues, including epithelial, fibroblast, myeloid, lymphoid and endothelial cells, collectively representing most cell types within the breast cancer TME (Fig. 1e-f; Extended Data Fig. 1b; Supplementary Table 1).

**Fig. 1.**
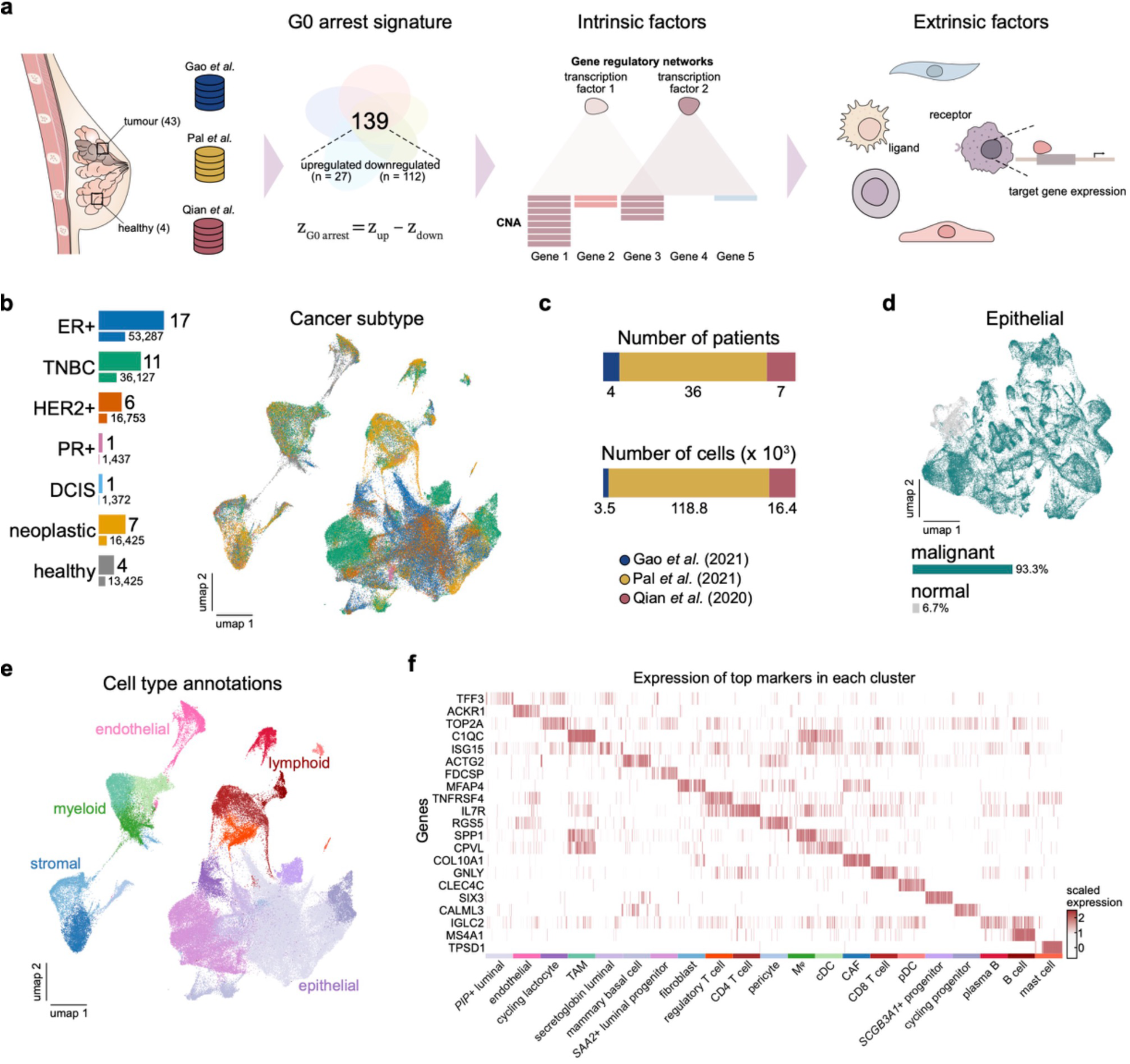
An overview of the single cell breast cancer compendium and analysis steps. **a** Schematic representation of the methodology employed in this study to analyse G0 arrest and proliferation states in single cell datasets. **b** Breast cancer subtypes are illustrated with distinct colours, where thicker bars (top) indicate the number of patients per subtype and thinner bars (bottom) represent the number of cells per tumour type. **c** Summary of the number of patients and corresponding cell counts in each study cohort. **d** Cell labels denoting malignant and normal epithelial cells inferred using inferCNV analysis. **e** UMAP visualisation displays the identified cell classes along with detailed cell type annotations. **f** Heat map of the top highly expressed genes per cell type, with annotations colour-coded to correspond with those in panel **e**.

We used inferCNV^41^ to identify malignant cells (Fig. 1d), harbouring a total of 432 copy number alterations, including amplifications in chromosomes 1q, 8q and 16p, as well as deletions in 3p,16q and 17p (Extended Data Fig. 1d). Genes affected by these alterations included known oncogenes^18^ such as *ERBB2* and *MYC*, and tumour suppressors *BRCA1* and *BRCA2*.

We used our previously developed G0 arrest scoring method^36^ to assess the proliferative capacity and cell cycle arrest frequency in malignant single cells (Extended Data Fig. 2a). To capture both extreme and intermediate phenotypes of proliferation and cell cycle arrest, we used stringent cut-offs defining three populations: fast-cycling (12.5%), slow-cycling (87.5%) and G0 arrested (12.5%) cells (see Methods, Fig. 2a; Supplementary Table 2). The differential abundance test demonstrated that the G0 arrested and fast-cycling labels of the tumour cells were indeed distinct cellular states (Extended Data Fig. 2b-c), confirming the two extreme proliferation capacities of these cells.

**Fig. 2.**
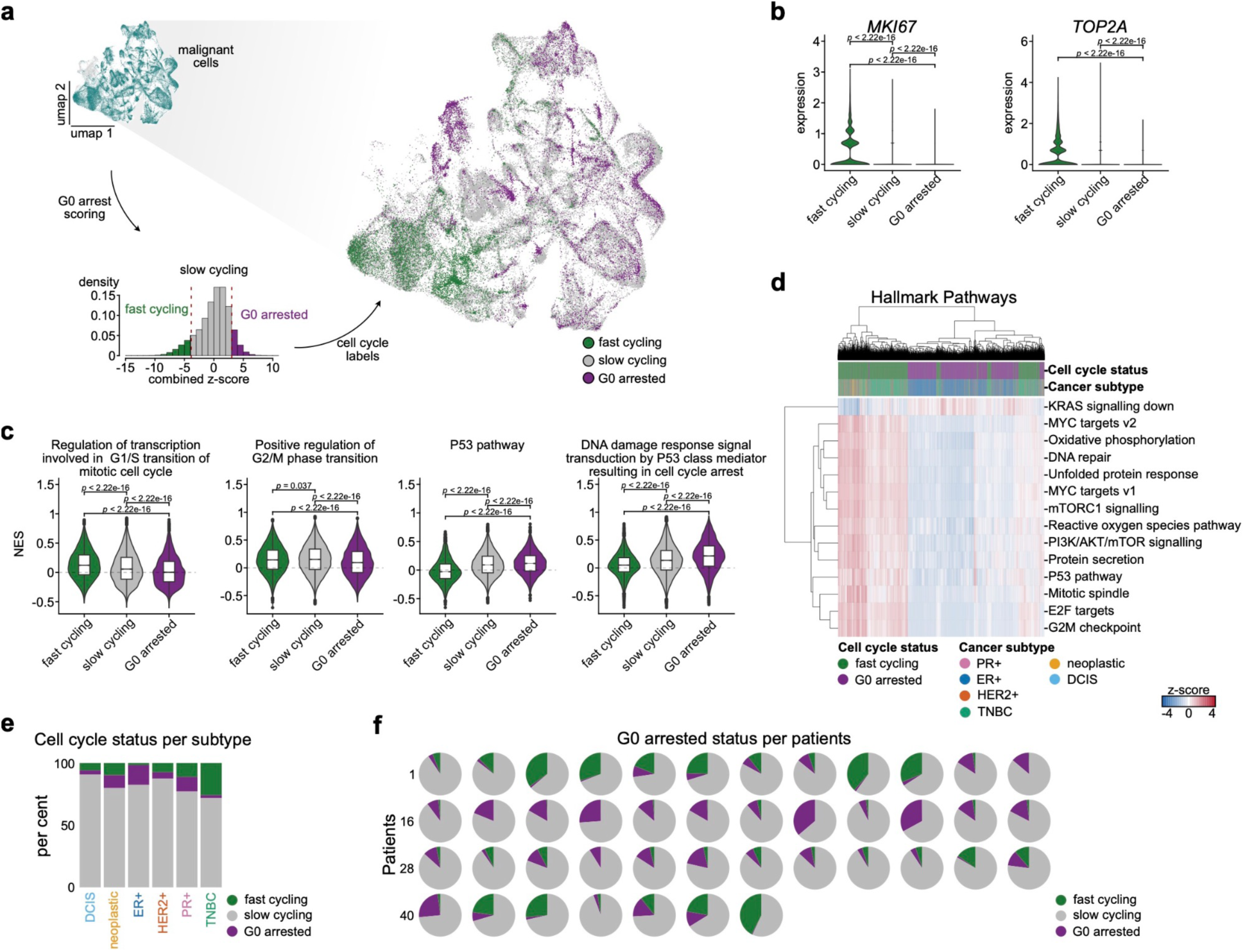
G0 arrest and fast cycling phenotypes in primary breast cancer. **a** Workflow for calculating G0 arrest scores, with UMAP embedding showing malignant cells annotated by discrete cell cycle states. **b** Expression comparison of proliferation markers *MKI67* and *TOP2A* across different cell cycle states. Wilcoxon rank-sum test p-values are indicated on the plots. **c** Gene Ontology term comparison across G0 arrested, fast-cycling, and slow-cycling states, with dots representing outliers, boxes extending to the 25th and 75th percentiles, and the median normalised enrichment scores (NES) displayed. P-value calculated using Kruskal-Wallis test. **d** Hallmark pathway enrichment analysis contrasting G0 arrested and fast-cycling cells. **e-f** Percentage of fast-cycling, slow-cycling and G0 arrested cells across breast cancer subtypes **(e)** and individual patients **(f)**.

Fast-cycling cells upregulated conventional proliferation markers (*MKI67*, *TOP2A*), cell cycle checkpoints, MYC/E2F targets and other pathways linked with cell proliferation in (Fig. 2b-d). In contrast, G0 arrested cells demonstrated p53-sustained DNA damage response (Fig. 2c). Thus, the distinct cell cycle states captured within these primary breast tumours bear the expected hallmarks of proliferation and quiescence, providing confidence that we had captured these switches. We also ensured that our G0 scoring method did not mistakenly capture low-quality cells^42^, as indicated by *MALAT1* expression (Extended Data Fig. 2d), nor a senescent state, as all malignant cells lacked senescence markers such as β-galactosidase (Extended Data Fig. 2e).

One would expect the prevalence of these cell cycle states to align with the classical breast cancer molecular subtypes. Indeed, the ER+ tumours presented the highest fraction of G0 arrested cells, whilst TNBC samples were abundant in fast-cycling cells (Fig. 2e-f), consistent with the clinical features of these subtypes^18,19^. Importantly, although G0 arrested cells were sometimes a minority, they were nevertheless present across all molecularly defined subtypes (Fig. 2e), highlighting this state is widespread in primary tumours and likely mediates dynamic cell adaptation. Patients with fast proliferating ER+ tumours had shorter relapse-free survival, whereas those with higher G0 arrest levels manifested longer latency before relapse (Extended Data Fig. 2f), resembling the indolent phenotypes driving metastatic latency described by Montagner et al.^43^. No significant survival differences were observed in TNBC (Extended Data Fig. 2g), indicating that our cell cycle scoring is not simply recapitulating the molecular subtypes of breast cancer but rather an independent clinically relevant feature, particularly in ER+ tumours.

### Genomic drivers of proliferation/G0 arrest capacity in single cells and large cancer cohorts

Unsurprisingly, G0 arrest was prevalent in primary tumour cells, especially within ER+ breast cancers. Cancer cells often exit the cell cycle as an adaptive response to stress during tumour development. While many G0 arrested cells in our cohort likely represent a transient, short-lived state reflecting rapid adaptation, others may be trapped in a deeper arrest, akin to dormant precursors associated with late recurrence. To interrogate which of these states were predominantly captured in our data, we reasoned that the genetic makeup of the proliferating and G0 arrested cells, reflecting their evolutionary distance, could also reflect the timing of G0 arrest: if we are simply capturing a short-lived cell cycle arrest state, the genomic profiles of these cells should resemble those of their proliferating counterparts; conversely, locking cells in a longer-lived G0 state would pre-empt the accumulation of further genomic alterations which may be observed in the fast-cycling cells. To determine the nature of these states, we examined the genomic profiles of proliferating and G0 arrested cells, as determined by inferCNV. Surprisingly, G0 arrested cells displayed fewer global CNA alterations, especially in ER+ patients (Fig. 3a-b, Extended Data Fig. 3), suggesting these cells may be “*stuck*” at an earlier tumour evolutionary stage, before the acquisition of several additional changes which can be observed in proliferating cells. It is therefore possible these cells have been in cell cycle arrest for longer than would be expected during a transient stress response. This aligns with our evidence that the G0 arrest score reflects the depth of cell cycle arrest^36^. Furthermore, the G0 cells also presented elevated markers of dormancy programmes previously reported in breast cancer (Extended Data Fig. 2h). Hence, by selecting the extreme G0 arrest and proliferation phenotypes, we have naturally enriched for a population exhibiting longer-lived cell cycle arrest.

**Fig. 3.**
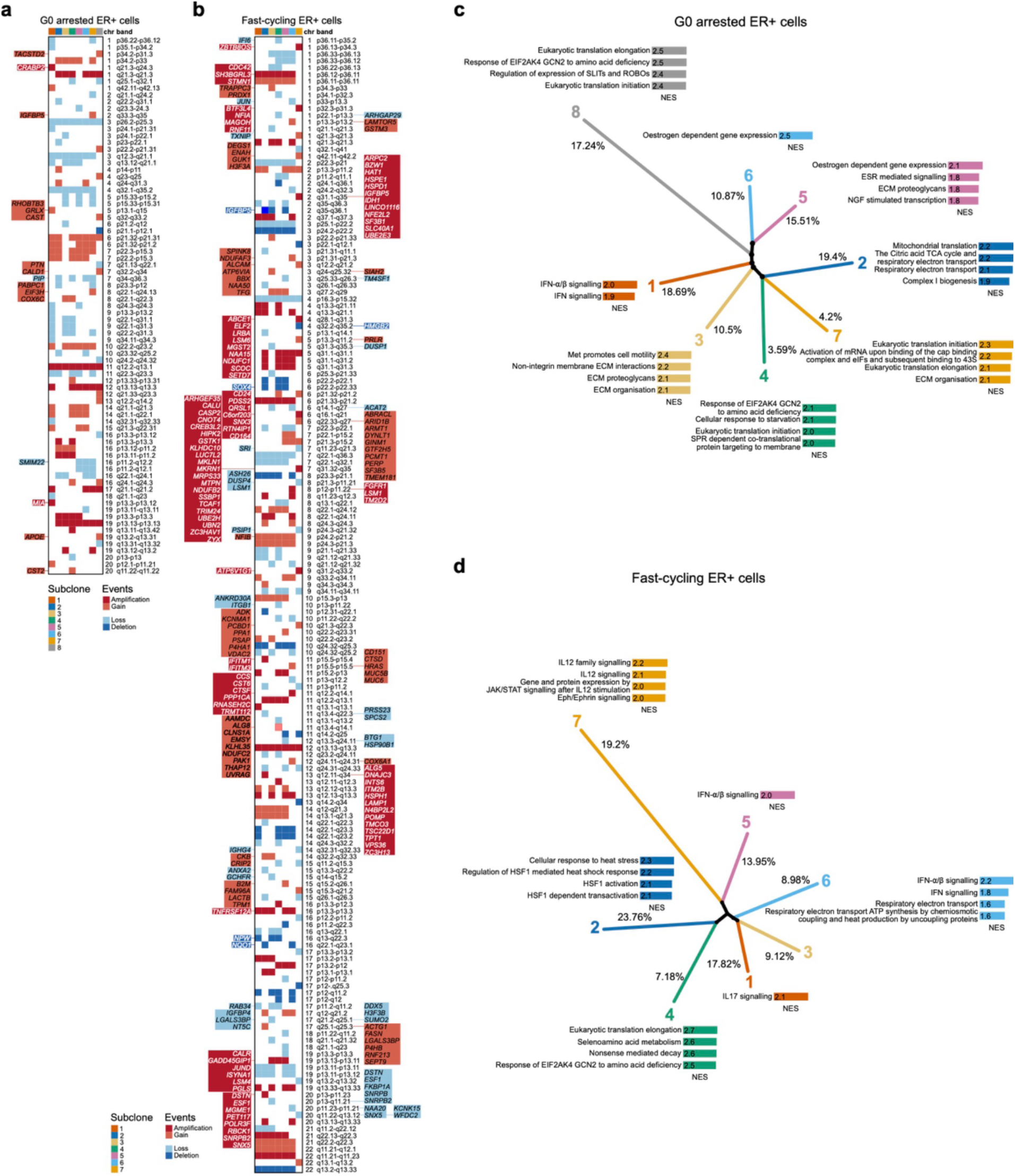
Genomic differences between G0 arrested and fast-proliferating ER+ breast tumours. **a** Chromosomal arm-level alterations in G0 arrested ER+ subclones, highlighting shared and unique genomic changes between subclones, as inferred by SCEVAN. Differentially expressed genes between G0 arrested and fast-cycling cells at each chromosomal location are highlighted on the side. **b** Chromosomal arm-level alterations in fast-proliferating ER+ subclones, illustrating key genomic differences compared to G0 arrested states. **c** Clone tree of G0 arrested ER+ tumour cells, with subclones colour-coded according to Reactome pathway enrichment of the correspondingly altered genes. **d** Clone tree of fast-proliferating ER+ tumour cells, with subclones colour-coded by Reactome pathway enrichment.

Copy number analysis in G0 arrested ER+ cells identified eight subclones with shared features, including losses on chromosome 3p (p26.2-p25.3) and amplification of 11q (q12.2-q13.1), linked with poor prognosis^44^ (Fig. 3a). Subclone 2 (19% of cells) was enriched for mitochondrial respiration changes (Fig. 3c), consistent with reduced energy demands of G0 arrest^45^. We also noticed copy-number driven response to starvation and metabolic stress (subclone 4), as well as subclonally altered translation and angiogenesis-regulating *SLIT*/*ROBO* pathways (subclone 8), capturing other features commonly linked with quiescence. Subclone 1 showed enrichment in interferon alpha and beta signalling, linked with chemotherapy resistance in inflammatory breast cancer^46^, along with amplifications in *CRABP2* and *MIA* genes implicated in epithelial-mesenchymal transition (EMT) and metastasis^47,48^. Other subclones (3, 5 and 7) were linked with cell motility and extracellular matrix remodelling, suggesting the acquisition of pre-migratory features.

Fast cycling ER+ cells also exhibited interferon signalling (subclones 5 and 6), and metabolic changes (subclone 4) but differed in dominant features including HSF1-mediated stress response (subclone 2, 24% of cells) and inflammation-driven CNAs (subclones 1 and 7, 37% of cells), suggestive of a genome-mediated adaptation in response to an immunogenic environment^49,50^ (Fig. 3d). TNBC showed similar genomic adaptations (Extended Data Fig. 3), with G0 cells exhibiting additional alterations in glucose metabolism and keratinisation (subclone 5), and fast-cycling cells showing interleukin signalling changes (subclone 1).

Gene-level CNA analysis highlighted known enablers of proliferation in fast-cycling cells, including *TP53* deletions, *MYC*, *PTEN* and *EGFR* amplifications^51,52^ (Extended Data Fig. 4a). Notably, the histone acetyltransferase *CREBBP*, a tumour suppressor in TNBC that drives aggressive growth when lost^53^, was deleted in fast-cycling cells but amplified in G0 cells, reflecting a potential genomic switch contributing to divergent proliferation/arrest phenotypes. The chromatin modulator *ASXL2,* promoting proliferation in ER+ breast cancer through ERα activation^54^, was amplified in fast-cycling cells and deleted in slower proliferating tumours from TCGA (Extended Data Fig. 4b). The most prevalent alteration in G0 cells was *ZFHX3* loss, associated with reduced proliferation in ER+ breast cancer^55^ and prostate cancer^56^ and EMT induction in mammary epithelial cell lines^57^. Intriguingly, *ZFHX3* deletion also ranked among top alterations in fast-growing metastatic tumours in the MET500 cohort but not in primary tumours from TCGA (Extended Data Fig. 4b-c). Thus, *ZFHX3* may mark the precursors of successful metastatic colonisation.

In conclusion, key features distinguishing proliferating from G0 cells appear selectively maintained through CNAs during cancer evolution. This suggests genetics may partly drive phenotypic plasticity at the extremes of cell cycle behaviour, mirroring the “*heritable plasticity*” described by Schiffman et al.^58^.

### Master regulators of proliferation-arrest switches in breast cancer cells

Beyond genomic alterations, gene regulatory networks (GRNs) can offer insights into tumour cell cycle regulation. We hypothesised that GRNs in G0 arrested cells could reveal conserved and subtype-specific transcription factors (TFs) driving cell cycle arrest. Using pySCENIC^59^, we identified six TF modules governing G0 arrest in invasive breast tumours (Fig. 4; Extended Data Fig. 5). These modules regulate cell cycle arrest (M1-M2), anti-tumour immunity via JAK-STAT signalling and interferon stimulation^60^ (M3), integrated stress responses (ISR) through ATF3, ATF4 and DDIT3 (M4)^61^, and metastasis-related processes such as EMT (M5) (Fig. 4a-b). Using our previously developed methodology to infer EMT macro-states in single cell data^62^, we were able to determine that these cells occupied a hybrid E/M state (Fig 4c). Additionally, a SOX9-regulated stemness module (M6) was also observed, which may manifest in a minor subpopulation of G0 arrested TNBC cells (Fig 4a-b).

**Fig. 4.**
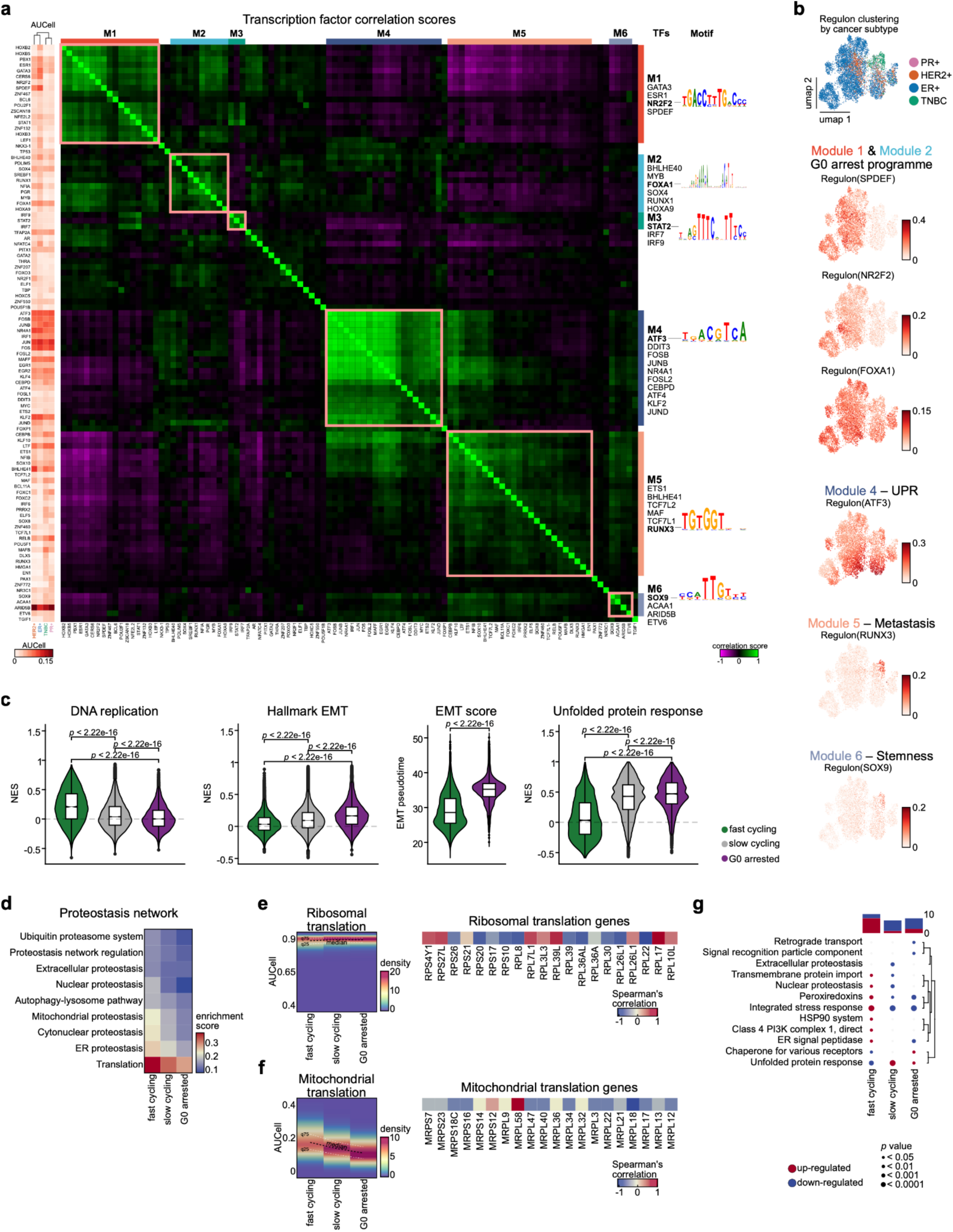
Gene regulatory networks modulating G0 arrest. **a** The left heat map shows AUCell enrichment scores for each regulon across breast cancer subtypes, as inferred by SCENIC. Middle heat map displaying correlation scores of the top 100 transcription factors (TFs) in G0 arrested cells. Vertical (right) and horizontal (top) bars highlight modules (M1–M6) with corresponding TFs, along with motifs for selected TFs. Inferred binding motifs enriched in selected TFs are shown on the right. **b** UMAP plot of G0 arrested cells (left) with regulon activities of selected TFs, categorised into biological functions. **c** Normalised enrichment scores (NES) for DNA replication, unfolded protein response, and hallmark EMT pathways, assessed by cell cycle status, and EMT score distribution by cell cycle status; dots indicate outliers, with boxes extending from the 25th to the 75th percentiles and the median indicated. Statistical significance (*p* value) was calculated using the Kruskal-Wallis test. **d** Heat map showing the enrichment of proteostasis pathways based on cell cycle state. **e-f** Ribosomal **(e)** and mitochondrial **(f)** translation enrichment, illustrating pathway activity depending on the cell cycle state (left), along with a breakdown of gene-level activity across the pathway (right). The gene-level activity is calculated as the Spearman correlation between the expression of the gene and the pathway score. **g** Upregulated and downregulated proteostasis pathways across cell cycle states.The bar charts on the top reflect the number of pathways up- and downregulated for each category, in red and blue, respectively.

Fast-cycling cells showed five GRN modules linked to proliferation, biosynthesis (M1-M3), IFN type II signalling (M4), and stemness/dormancy regulated by SPDEF and NR2F1 (M5)^63^ (Extended Data Fig. 5a). *MYBL2* and *FOXM1* regulation were enriched across molecular subtypes (Extended Data Fig. 5b).

Subtype-specific regulons highlighted distinct cell cycle control. In ER+ tumours, G0 arrest was most prominently regulated by classical luminal markers (*GATA3*, *FOXA1*). TNBC tumours displayed higher regulon specificity scores of *MYC* and *RUNX3*, well-known proliferation regulators (Extended Data Fig. 5d). Notably, PR+ cells exhibited stem-like characteristics in both fast cycling (*SOX6*, *SOX8*) and arrested cells (*POU5F1*/*OCT4*), whereas cell fate commitment factors (*SOX2*, *SPDEF*) predominantly affected fast cycling cells in HER2+ cancers.

Slow-cycling cells activated GRNs shared by proliferating and arrested cells, regulating proliferation, transcription, cell fate commitment and ISR. These cells displayed significant activity of the *BCL11A*, *ATF3* and *JUN* regulons (Extended Data Fig. 5e-f). Their hybrid characteristics suggest a transitional state, enabling rapid shifts between proliferation and arrest to adapt to the tumour niche.

### Proteostasis pathways govern stress adaptation and G0 arrest persistence in breast cancer

Proteostasis pathways are critical for maintaining protein folding, degradation and overall proteome integrity, enabling cells to adapt to stress and to balance G0 arrest and proliferation^64^. Our GRN analysis highlighted the importance of proteostasis mechanisms, such as the UPR (Fig. 4c), across cell cycle states. We observed a gradient of reduced expression of translation, endoplasmic reticulum (ER) proteostasis and mitochondrial proteostasis genes correlating with cell cycle rates, with the lowest activity at G0 (Fig. 4d). While ribosomal translation remained active across all malignant cells, mitochondrial translation showed a declining trend (Fig. 4e-f), with most related genes negatively correlated with cell cycle arrest (Fig. 4f), reflecting the lower energy demands of G0 cells.

Further breakdown of proteostasis pathways unveiled an upregulation of HSP90 molecules, PI3K complexes, ER signal peptidases and peroxiredoxins in fast proliferating cancer cells (Fig. 4g). In contrast, G0 arrested cells downregulated retrograde transport and signal recognition particles, whereas slow-cycling cells showed a downregulation of extracellular and nuclear proteostasis. UPR activity increased in both G0 arrested and slow-cycling cells, though ISR genes were largely downregulated (Fig. 4g). This aligns with reports that in addition to promoting gene expression, ATF4 and ATF3 can act as transcriptional repressors under specific stresses to regulate glucose metabolism and lipid homeostasis^65–67^. Overall, this analysis demonstrates the remarkable capacity of G0 arrested cells to remodel their proteostasis network to achieve a “*slow flow*” state of reduced but viable activity within the tumour.

### Cell cycle state-dependent rewiring of tumour cell-TME interactions in single cell data

Given that the TME influences the induction and maintenance of growth arrest^68^, we next interrogated whether tumour-TME interactions are rewired when cells switch between proliferation and cell cycle arrest. First, we employed NicheNet 2.0^69^ to explore the influence of various ligands expressed by immune and stromal cells on cancer cell cycle decisions. *TGFB1* emerged as the top ligand in G0 arrested cells, driving expression of cell cycle checkpoint regulators (*EGR1*, *FOS*, *KLF6*) and suppressing cyclin-dependent kinase activity (Fig. 5a, Extended Data Fig. 6a-b), reinforcing the known role of integrin-activated TGF-β1 in inducing cell cycle arrest^70,71^. Notably, *TGFB1*-induced expression of *MGP* in G0 arrested cells may enhance chemoresistance via extracellular matrix interactions^72^. Additionally, Oncostatin M (OSM)-induced *TIMP1* expression could disrupt cell cycle dynamics by modulating *CDK1* and *CDKN1A*^73,74^. Stromal ligands, including *FGF1*, *TGFB3*, *IGF1* and *TNF*, strongly correlated with genes regulating all cell cycle phases in fast-cycling cells (Extended Data Fig. 6c-d; Supplementary Table 3). Notably, T cell-specific interferon gamma (*IFNG*) elevated *HLA-A* expression in these cells (Extended Data Fig. 6c-d).

**Fig. 5.**
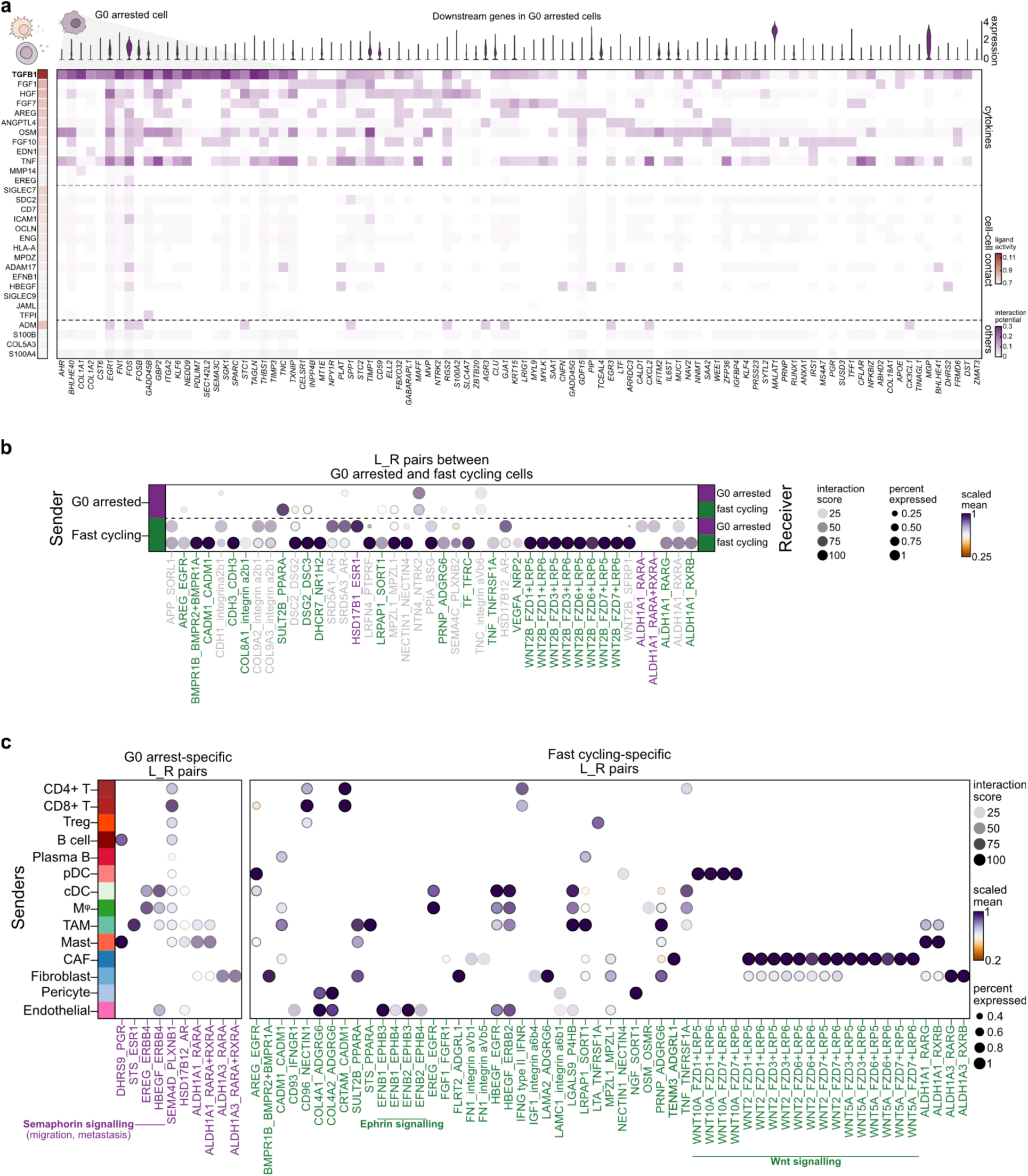
Cell cycle state-specific interactions with the TME. **a** The top 30 ligands expressed by cells in the TME and their corresponding targets in G0 arrested cancer cells, as inferred by NicheNet. Rows highlight ligands on immune and stromal cells, alongside their activity depicted by a white-red colour gradient. The ligands are grouped manually into biologically relevant categories, annotated to the right. Columns depict downstream targets in G0 arrested cancer cells. Violin plots (top) display the expression levels of these target genes. **b** Ligand-receptor interactions between G0 arrested and fast-cycling cancer cells, as inferred by CellPhoneDB. Sender cells are shown on the left, receptor cells are shown on the right. The purple ligand-receptor pairs indicate interactions in which G0 arrested cells function solely as receivers, regardless of the origin of the sender. The green ligand-receptor pairs denote interactions where fast-cycling cells exclusively serve as receivers, irrespective of the origin of the sender. The grey ligand-receptor pairs mark interactions that are not unique in either category of cell types. **c** Ligand-receptor interactions established uniquely between cells in the TME and G0 arrested cells (left panel) or fast-cycling cells (right panel), illustrating distinct communication networks depending on the cell cycle state. The rows depict the sender cells in the TME, the columns highlight distinct ligand-receptor pairs with the receiver tumour cell, as inferred by CellPhoneDB.

Using CellPhoneDB^75^, we assessed ligand-receptor interactions across different cells in the TME. G0 arrested cells showed reduced communication with other cells, including proliferating or arrested tumour cells (Extended Data Fig. 6e, Figure 5b). However, semaphorin-plexin signalling, known to promote EMT and metastasis^76,77^, was uniquely observed between G0 cells and various immune cells, including CD8 T cells and tumour-associated macrophages (TAMs) (Fig. 5c). TAMs and DCs/macrophages also influenced hormone signalling (STS-ESR1) and ERBB4-driven growth pathways in G0 cells, respectively (Fig. 5c; Supplementary Table 3). Conversely, in fast-cycling cells we observed CAF-induced activation of the Wnt pathway alongside myeloid stimulation of *EGFR* and *ERBB2* (*HER2*), all known to drive proliferation^78^, endothelial-linked Ephrin signalling, implicated in cell adhesion and migration^79^, and increased communication with CD4/CD8 T cells (Fig. 5c). The *LGALS9*-*P4HB* pairing between fast-cycling cells and TAMs has been linked with immunosuppression^80^.

Retinoic acid signalling appeared in both G0 (*ALDH1A3*-*RARA*) and fast-cycling (*ALDH1A1*-*RXRB*, *ALDH1A3*-*RARG*) cells, though its role may vary, supporting stemness maintenance^24^ in G0 and proliferation via the non-canonical Wnt^81^ in fast-cycling cells (Fig. 5c). Shared interactions involving stromal collagens, *TGFB3* and tumour cell-specific integrins (α2β1, αVβ6, αVβ8), mostly mediating interactions with CAFs, may regulate ECM organisation in both states (Extended Data Fig. 6e-f).

To assess the potential for CD8 T cell exhaustion, we analysed naive, effector memory, pre-exhausted, exhausted and short-lived exhausted cell (SLEC) subsets (Extended Data Fig. 7a-d). In line with our previous observations of increased immune reactivity, fast-cycling cells presented more interactions with exhausted T cells than G0 cells (Extended Data Fig. 7f), mediated by cytokines like *ANGPTL4*, shown to reprogram CD8 T cell metabolism^82^ (Extended Data Fig. 7e), and *TIGIT*-*NECTIN2* signalling (Extended Data Fig. 7f). However, T cell exhaustion appeared more strongly linked with DCs and TAMs (Extended Data Fig. 7f).

### Spatial organisation of proliferation and G0 arrest phenotypes unveils extensive niche remodelling

While single-cell data offer insights into the intrinsic regulation of proliferation and cell cycle arrest in cancer cells, the spatial emergence of these states within tumour tissue remains unclear, despite being essential for understanding how these dynamics drive tumour growth. To investigate this, we examined G0 arrest and proliferation phenotypes using spatial transcriptomics data from 12 breast cancer 10x Visium slides (see Methods). Using our SpottedPy method^83^, we mapped G0 arrest and fast-cycling tumour hotspots and spatial interactions within the TME (first panel in Fig. 6a, Extended Data Fig. 8a). Figure 6a illustrates the spatial organisation of these hotspots in two patient tissues (additional patients in Extended Data Fig. 8a). While most of the tissue was composed of tumour cells with intermediate cycling capacity, the extreme proliferation or cell cycle arrest areas were dispersed throughout the tissue rather than collocated in a single region, suggestive of the tumour’s ability to fine-tune proliferation rates locally to adapt to varying stressors.

**Fig. 6.**
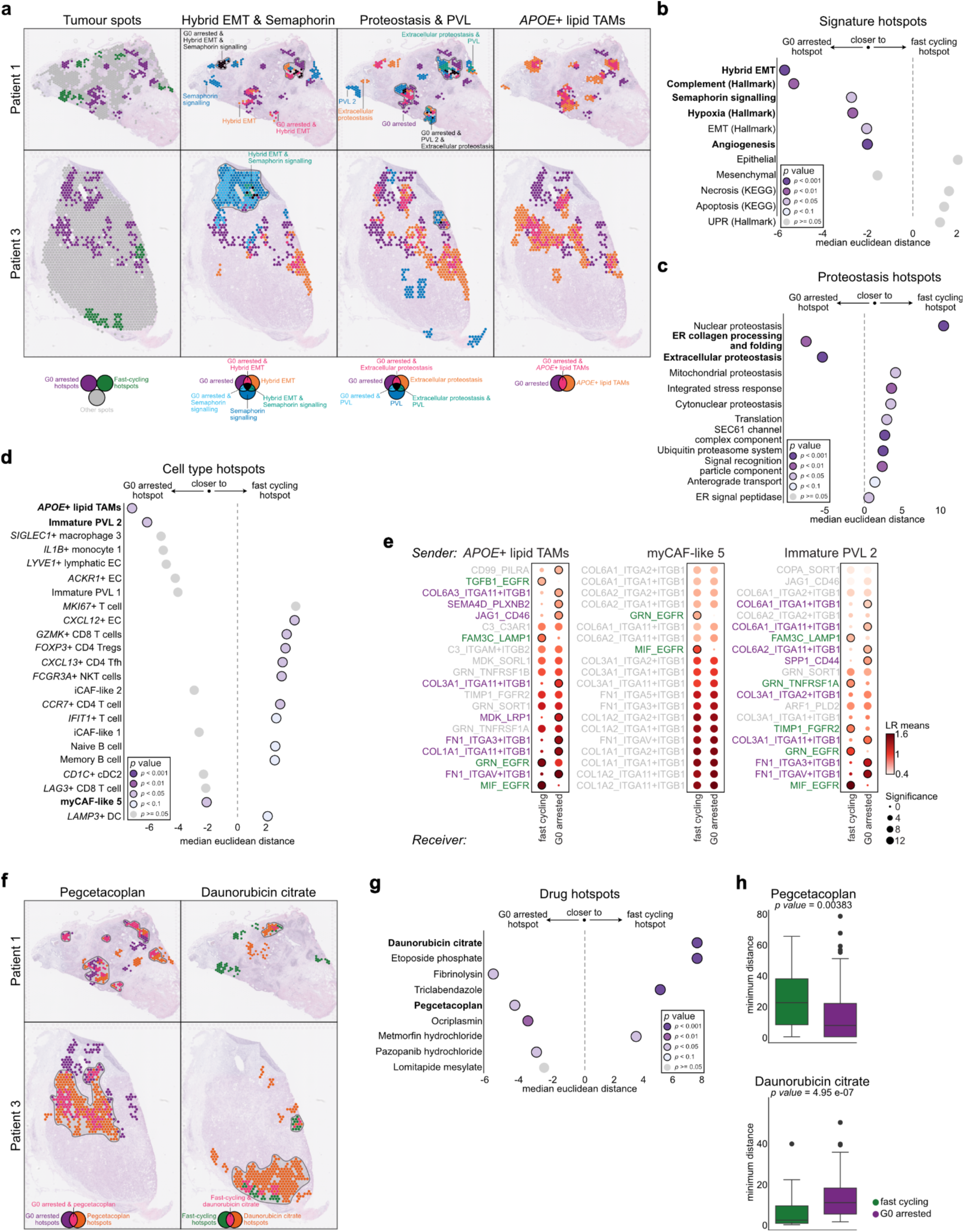
Spatial organisation of G0 arrest and high proliferation hotspots in breast cancer. **a** Spatial transcriptomics images showing hotspots identified by SpottedPy in two patient samples. First panel: Hotspots are colour-coded by cell cycle state: green (fast-cycling) and purple (G0 arrested), while grey marks the remaining tumour spots. Second panel: Overlap between G0 arrested tumour hotspots, hybrid EMT hotspots and semaphorin signalling hotspots. Third panel: Overlap between G0 arrest hotspots, proteostasis signalling and immature perivascular-like cell (PVL) hotspots. Fourth panel: Overlap between G0 arrest hotspots and *APOE*+ lipid-associated TAMs. **b-d** Bubble plots representing relative median Euclidean distances to G0 arrest cancer hotspots (left) or fast cycling hotspots (right) for **(b)** selected cancer hallmarks and pathways, **(c)** proteostasis pathways and **(d)** immune and stromal cell hotspots. Circles with a black contour depict significant p-values (i.e. hotspots that are significantly closer to either G0 or fast cycling pockets). **e** Ligand-receptor interaction analysis between *APOE*+ lipid-associated TAMs, myCAFs and immature PVL hotspots as senders and tumour cells as receivers, in relation to G0 arrested and fast-cycling hotspots. Purple indicates ligand-receptor interactions unique to G0 arrested tumour cells, green indicates those unique to fast cycling cells. **f** Drug-niche hotspots identified in the same two patients from a, showcasing areas within the tumour where the drug is predicted to be most effective and their overlap with G0 arrest cancer hotspots (left, shown for pegcetacoplan) or fast cycling hotspots (right, shown for daunorubicin citrate). **g** Distances between drug-effective regions and either G0 arrest or fast-cycling hotspots. The drugs shown were found to be differentially associated with G0 arrest or fast cycling areas within the tumour by drug2cell. **h** Box plots depicting tumour-drug area distances for the two drugs most strongly correlated with G0 arrest and fast-cycling niches.

We next analysed the spatial proximity of G0 arrest/fast-cycling hotspots and overlap with key signatures, including EMT, hypoxia, angiogenesis, proteostasis, apoptosis and various cell types within the TME. We found that tumour spots in a hybrid E/M state^62^ (and to a lesser extent a fully mesenchymal state) were significantly closer to G0 arrest hotspots (Fig. 6a second panel, 6b), although some intra- and inter-patient variability could be observed (Extended Data Fig. 8b). These observations suggest an interplay between G0 arrest and E/M transformation akin to that observed in the single cell data. In addition, G0 arrest hotspots tended to be proximal to pockets of hypoxia and angiogenesis, but lacked apoptosis hallmarks, indicating they were not sites of programmed cell death (Fig. 6b, Extended Data Fig. 8b). Furthermore, we revisited the proteostasis pathways observed in single-cell data and found collagen processing and folding in the ER, as well as extracellular proteostasis activity in the vicinity of G0 arrest hotspots (Fig 6a-c). Given that mediating collagen production is a primary function of the UPR, our data suggest that G0 cells remodel their proteostasis network to influence not only intrinsic proteome homeostasis, but also the structure/composition of their surrounding environment^81^.

Interestingly, G0 arrest hotspots appeared encapsulated within *APOE*+ lipid-associated TAM pockets and were near immature perivascular cells (PVL) and myofibroblastic cancer-associated fibroblasts (myCAFs), while remaining distant from lymphoid cells (T, NK and B cells) and monocytes (Fig. 6a,d). This pattern suggests a tumour protective niche that may specifically enable the immune evasion of G0 arrested cancer cells, while the fast-proliferating areas are more prone to cytotoxic T cell recognition. We noted enriched *SEMA4D*-*PLXNB2* interactions in the neighbourhood of several G0 arrest hotspots, corroborating the modulatory role of G0 states by semaphorin signalling observed in single-cell data (Fig. 6a-b, e). Further spatial ligand-receptor analysis between *APOE*+ lipid-associated TAMs and G0 arrested cells (Fig. 6e) confirmed enriched *SEMA4D*-*PLNXB2* pairing (Fig. 6e, Extended Data Fig. 9a). Plexin B2 interactions with semaphorin IV have been implicated in tumour invasion through cytoskeletal reorganisation^84,85^ and have also been reported by Borrelli et al.^86^ to play a key role in metastatic colonisation by triggering an epithelialization switch in the metastatic precursor cells as a requirement for adaptation in the new environment. As our analysis suggests an EMT rather than mesenchymal-epithelial transition (MET) activation in these areas, this could imply a pleiotropic role for semaphorin IV-plexin B2 signalling capable of toggling between EMT and MET depending on the context (primary tumour/metastasis). Therefore, the G0 arrest hotspots display some characteristics akin to those of seeder cells of metastases, suggestive of areas that are actively adapting to microenvironmental pressures.

We also found the *JAG1*-*CD46* ligand-receptor pair enriched, indicating a potential activation of the complement response in the G0 arrested cells (Fig. 6e). Indeed, significant activity of the complement pathway was detected in the proximity of G0 arrest hotspots (Fig. 6b). In addition, collagen/integrin signalling, particularly between the collagen 3/6 families and integrin ɑ11β1, was prevalent in mediating interactions between G0 arrested cells, *APOE*+ lipid-associated TAMs, myCAFs and immature perivascular cells (Fig 6b, Extended Data Fig. 9). Integrin ɑ11β1 has been shown to regulate CAF stiffness and promote metastatic potential in non-small cell lung cancer^87^. In conjunction with the enriched extracellular proteostasis and collagen processing signals observed proximal to the G0 cancer niche (Fig. 6c), these analyses suggest that the niche may be extensively remodelled, both through a reorganisation of the extracellular matrix and of the vasculature surrounding the G0 cancer cells. This may be enabled through the co-option of *APOE*+ lipid-associated TAMs and myCAFs which may be secreting factors to enable the remodelling. Another tantalising hypothesis is that the G0 cancer cells could be secreting some of these factors, as shown in our previous work^87^ reporting multiple collagen and integrin molecules to be potentially secreted by quiescent cells. The extensive intrinsic and extrinsic remodelling of G0 cells and their immediate environment may promote intravasation into the neighbouring blood vessels, which are likely being actively expanded as suggested by the presence of immature perivascular cells.

### Segregation of G0 arrest and proliferative areas suggests differential responses to drugs

The observed spatial segregation between G0 arrest and proliferative areas might inform therapy response. Using drug2cell^88^, which predicts targets of drugs from the ChEMBL database based on their single cell expression, we identified molecular hotspots with distinct drug sensitivities. This analysis revealed that proliferative niches prominently overlapped with daunorubicin citrate-responsive regions (Fig. 6f). Daunorubicin, a DNA replication inhibitor commonly used in leukaemia, targets vascular mimicry structures–an adaptive mechanism supporting tumour growth and survival. Its ability to suppress mediators such as *TGFB1* and *MMP2*^89^ has potential utility in breast cancer, particularly when combined with tamoxifen to eliminate both cancer cells and stem cells^90^. In contrast, G0 arrest hotspots were associated with potential efficacy for pegcetacoplan, a complement pathway C5 inhibitor. This adds to our earlier findings that G0 niches display reduced complement pathway activity (Fig. 6e). Pegcetacoplan’s role in modulating inflammation^91^ suggests it could counter immune evasion mechanisms within quiescent tumour niches. These findings underscore the spatial heterogeneity of drug-niche interactions and propose potential combination therapies targeting different tumour compartments.

In summary, our analysis delineates distinct quiescent, potentially pre-migratory niches within primary tumours, rich in tumour-promoting signals associated with local stress adaptation. Surprisingly, these niches appear to be dispersed across the tumour, rather than clustering at the leading edge, hinting at proliferative and G0-hybrid E/M patterns that fluctuate throughout the tissue as a response to intrinsic and extrinsic stress factors.

### A foundation model for G0 arrest and proliferation in single cells

To generalise the identification of G0 arrest, slow-cycling, and fast-cycling states in other single-cell datasets, we employed a foundation model framework similar to ones we have developed previously^92^ to train a classifier of these states (Extended Data Fig. 10a). The proposed single-cell Large Language Model (scLLM), G0-LM, demonstrated robust performance in classifying G0 arrested, slow-cycling and fast-cycling states, achieving an average AUROC score of 0.94 across these categories (Extended Data Fig. 10b), underscoring the model’s efficacy in distinguishing biologically distinct cell cycle states. Additionally, the embeddings generated by G0-LM displayed notable patterns consistent with biological expectations. Despite being trained solely for classification, the embeddings revealed a continuum, showing a gradual transition of cells from fast-cycling to slow-cycling and ultimately into the G0 arrest state (Extended Data Fig. 10c-d). This highlights the model’s capacity to capture progressive state changes. The ability of G0-LM, powered by scLLM, to delineate such trends suggests a high sensitivity to the complex gene expression landscapes associated with the G0 arrest phenotype. To enable future analyses of G0, slow and fast proliferation phenotypes in single cell breast cancer data, we provide the G0-LM model at https://github.com/secrierlab/G0-LM.

## Discussion

Our study reveals the interplay of cell cycle states within primary breast tumours, identifying proliferative and quiescent populations with distinct genetic and transcriptional profiles shaped by the TME. We observed a subtype-dependent prevalence of these states: ER+ tumours harboured a greater proportion of G0 arrested cells, while TNBC samples were rich in fast-cycling cells (Fig. 2d), reflecting known differences in clinical outcomes. G0 cells in ER+ tumours had fewer CNAs and expressed dormancy markers, suggesting they may be stably arrested at an earlier evolutionary stage rather than simply responding to transient stress. In contrast, fast proliferating cells upregulated the dormancy marker *NR2F1*, indicating the potential to develop a more agile, plastic and epigenetically-driven dormant state, not observed here and likely only activated by dormancy-promoting immune cells^61^ or upon treatment. Both cell states upregulated stemness modules, which may be decoupled from the strict cell cycle programme, although enacted through different transcription factors. The identification of a slow-cycling population bridging proliferative and quiescent characteristics suggests these cells could facilitate dynamic state shifts in response to TME pressures, enhancing adaptability.

G0 arrested cells were often in a hybrid E/M state, aligning with the “*go-or-grow*” model of cancer^27,93–95^. They also exhibited UPR and reduced mitochondrial translation, allowing them to remain in a viable but metabolically reduced state over prolonged periods. It has been shown that mitochondrial stress couples activation of the ISR with reduced mitochondrial translation^96,97^. This is highly consistent with our observations in G0 arrested cells, suggesting that mitochondrial remodelling and metabolic adaptation may be what underlies activation of ISR-associated TFs (i.e. ATF4, ATF3 and DDIT3/CHOP). Since cytosolic ribosomal gene levels remained stable, translation may be reduced through the activation of the ISR, allowing G0 arrested cells to be “*primed*” for an increase in protein synthesis in case exit from G0 is required^98^.

Our study also highlights the role of extrinsic factors in sustaining cell cycle arrest. Spatial profiling of breast tumours revealed patient-specific activity hotspots, identifying a G0 arrest cancer niche enriched in myCAFs and *APOE*+ lipid-associated TAMs, primed by semaphorin-plexin signalling and perivascular remodelling. Extracellular proteostasis signalling within these niches suggests active communication and remodelling of the ECM and TME through multiple collagen-integrin interactions. Due to imperfect resolution of the Visium data, the source of secreted signals remains unclear, but two scenarios are plausible: (1) the *APOE*+ lipid-associated TAMs and myCAFs may send signals, promoting an arrested, partially EMT-transformed state in tumour cells, or (2) tumour cells themselves may secrete signals to remodel the ECM and vasculature, which is developing with the help of the immature pericytes, and potentially co-opting TAMs and myCAFs to evade immune recognition. This spatial pattern suggests G0 arrested cells in primary breast tumours may already acquire traits enabling them to occupy pre-metastatic niches, with *APOE*+ lipid-associated TAMs and myCAFs priming these cells for migration and colonisation. These interactions sustain quiescence and remodel the niche, potentially facilitating intravasation into the nascent vasculature^86,99,100^. These findings highlight the potential of therapies targeting cancer-TME interactions to destabilise quiescent niches, particularly in ER+ tumours with high proportions of G0 arrested cells.

While our findings align partly with Baldominos et al.^23^, who noted a hypoxic quiescent niche containing immune-suppressive fibroblasts and exhausted T cells in TNBC tumours, we did not observe T cells near G0 hotspots. This is likely because Baldominos et al.^23^. specifically selected T cell-resistant tumour cells, which would be more likely to be rarer quiescent cells situated within the cytotoxic and exhausted proliferative hotspots observed in our analyses. Thus, our analysis highlights a different G0 arrest phenotype within the developing breast tumour that is likely not simply epigenetically driven as a reaction to T cell attack.

In our single cell and spatial analyses, we find some hints that G0 arrested cells may exhibit markers of a long-lived, dormant-like phenotype with hybrid E/M and tumour invasion/migration features. This indicates that our analysis captures not only rapid adaptation to changing microenvironmental stress but may also pinpoint tumour regions likely to harbour precursors of metastasis and recurrence. While not all G0 arrested cells become successful persister cells or metastatic seeders, our method offers a systematic “*first scan*” for potential pre-migratory areas in spatial data. However, fast proliferating tumours show worse outcomes in breast cancer, and Lucas et al.^101^ recently suggested that high-proliferating clones in single-cell DNA-sequencing datasets have increased metastatic potential in lung cancer. While Lucas et al.^101^ did not distinguish G0 cells as a separate category and metastatic potential may vary in breast cancer, we cannot rule out that dormancy-priming by *NR2F1* in fast-cycling cells may enhance their ability to switch into this programme under favourable microenvironmental conditions or therapy. This plasticity may outcompete the more constrained plasticity of metabolically inert G0 cells. The positioning of fast-cycling cells within cytotoxic and exhausted niches suggests that these areas may respond to immunotherapy, though this also emphasises the difficulty in eradicating the entire tumour, as immunotherapy is unlikely to be effective in G0 cancer pockets.

Together, these analyses lay the groundwork for future studies, once several limitations are addressed. Our definition of G0 arrest relies on a conservative threshold on gene expression scores, which could be adjusted to capture gradual state changes, especially in spatial data. However, the dynamic emergence and extinction of cancer niches cannot be inferred from a single time point. Future studies should use longitudinally profiled samples, ideally both from primary and matched metastatic tumours, to gain insights into how these niches evolve over time, their stability and potential to generate migratory precursors to metastasis. Secondly, while we used CNAs to estimate the relative timing of G0 arrest in relation to proliferative cells, we acknowledge that transcriptomic copy number estimation can be prone to error and susceptible to the reference employed. While identified genomic changes previously implicated in cell proliferation and breast cancer provide confidence in these findings, future studies should combine single-cell DNA/RNA sequencing with lineage tracing to draw more robust conclusions. Thirdly, this study has not explored epigenetic or metabolic adaptations required to stabilise G0 arrest. Although seminal studies such as by Rosano et al.^22^ noted that long-term quiescence is primarily driven by non-genetic adaptations to endocrine therapies, our analyses suggest that, at least in a pre-treatment setting, genomic alterations may predispose tumour cells to favour either fast proliferation or prolonged arrest. Further studies employing methylation, chromatin conformation and metabolomics are needed to explore the non-genetic mechanisms contributing to the evolutionary role of G0 arrest in cancer. We also acknowledge that the limited number of spatial transcriptomics samples may not fully capture the spatial variability in breast cancer, and the lack of single-cell resolution in the Visium platform complicates the inference of cell-specific signals, despite our deconvolution efforts. Lastly, while we identified intermediate (slow-cycling) cells as potential mediators of state transitions, further research is required to elucidate the molecular mechanisms behind these transitions and the TME factors involved.

In conclusion, our study advances the understanding of cell cycle regulation in breast tumours by revealing cell-intrinsic and extrinsic factors that shape proliferative dynamics. We identify a long-lived population of G0 arrested cancer cells with genetically-constrained transcriptional plasticity, alongside spatially organised niches characterised by immunosuppressive and cytotoxic microenvironments. These insights are particularly relevant for ER+ cancers, where targeting quiescent populations and disrupting dormancy-associated signalling networks, such as semaphorin-mediated pathways, may reduce recurrence risk. Additionally, our spatial mapping of proliferation and G0 arrest hotspots offers a foundation for tailoring therapies to the unique microenvironmental contexts of each niche.

## Methods

### Data collection, integration and cell type annotation

Single-cell RNA sequencing matrices for breast cancer, covering both non-invasive and invasive breast carcinoma samples, were sourced from the Curated Cancer Cell Atlas (https://www.weizmann.ac.il/sites/3CA/). These datasets underwent standardisation based on droplet-based studies conducted by Gao et al.^37^, Qian et al.^39^, and Pal et al.^38^.

To preprocess and integrate the samples, the standard Seurat pipeline with `SCTransform()` v2 normalisation^102^ was employed, wherein low-quality cells (with mitochondrial transcript percentage > 20) were removed and cell cycle phases were regressed out. Subsequently, the Leiden community detection algorithm was applied with a resolution of 1.2 on the 29 principal components of the integrated data, followed by uniform manifold approximation projection (UMAP) for dimension reduction.

Cell type annotation was performed at two resolutions. Initially, main cell types were identified using cell type signatures from PanglaoDB^103^ and *ScType* R package^104^. Then, for further refinement, the CELL type iDentification tool on the Deeply Integrated Human Single-Cell Omics platform^105^ was employed, by utilising averaged expressions of Leiden communities to accurately annotate cell types. Additionally, the top two differentially expressed genes of the annotated Leiden communities were calculated on the integrated data with a log2 fold-change of 0.25 and minimum percentage of 0.25 using the likelihood-ratio test for single-cell feature expression^106^.

To identify malignant cells within the epithelial cluster, the *inferCNV* R tool^41^ was utilised to infer copy number alterations (CNAs) from gene expression data. InferCNV was configured with the following parameters: a cutoff of 0.1, deemed suitable for 3’ sequencing droplet-based assays, `denoising = TRUE`, `HMM_type = “i6”` for Hidden Markov Model (HMM)-based CNA prediction, and `analysis_mode = “subclusters”` to identify clonal CNAs. A heat map depicting the relative expression intensities across each chromosome was generated, revealing regions of the tumour genome that were over- or under-abundant compared to normal cells.

### Differential gene expression analysis

We conducted differential gene expression analysis on the integrated dataset using Seurat’s `FindAllMarkers()` function on the RNA layer of the Seurat object with parameters `min.pct` set to 0.5 and `logfc.threshold` of 0.25 for only positively upregulated genes for both cell type annotations and cell cycle states.

### Quantification of G0 arrest in malignant single cells and spatial spots

First, based on inferred CNAs, the malignant cells were subsetted to implement a combined z-score, as previously defined by us in Wiecek et al.^36^. Z-scores of the 27 upregulated and 112 downregulated genes in the G0 arrest signature were then calculated using the GSVA R package. Next, a final G0 arrest score was obtained by subtracting the two scores. Finally, the G0 arrest scores were used for the discretisation of malignant cells into three categories: (1) G0 arrested, (2) fast-cycling and (3) slow-cycling. The cut off values of −3.83 and 3.03 were estimated by employing the *maxstat* R package on the G0 arrest scores, which utilises maximally selected rank statistics with several p-value approximation^107^.

The single-cell differential abundance testing was then performed on the cells with two extreme proliferating characteristics, and the neighbourhoods on a nearest neighbour graph were identified by employing the *miloR* R package^108^.

In spatial transcriptomics, z-scores for G0 arrest signatures within tumour spots were calculated similarly. Given the heterogeneity within spatial spots, we defined cut-off values using the 25^th^ and 75^th^ percentiles of G0 arrest scores. Tumour spots with z-scores below the 25^th^ percentile were classified as fast-cycling, while those above the 75^th^ percentile were classified as quiescent. This quantile-based approach accounted for the spatial variability inherent in tumour microenvironments and maintained consistency with the single-cell classification criteria.

### Genomic alterations in primary tumour cells

We used *SCEVAN* R package^109^ to investigate tumour clonality, Reactome Pathway enrichment of each subclone and oncoprint-like plots highlighting clone-specific alterations in fast-cycling and G0 arrested tumours.

### Genomic associations with G0 arrest/proliferation in single cells and large cancer cohorts

Gene-level copy number changes called by inferCNV in the single cell data were compared between G0 arrested and fast-cycling cell populations, and changes significantly enriched in either category were identified using a Fisher’s exact test, with Benjamini-Hochberg adjustment of p-values for multiple testing.

RNA sequencing data and gene-level copy number data from primary tumours from the TCGA Pan-Cancer Atlas were downloaded using *TCGAbiolinks* R package^110^. FPKM-normalised RNA-sequencing data and segmented copy number data from metastatic tumours were downloaded from the MAGI MET500 server (https://met500.med.umich.edu/downloadMet500DataSets), along with processed data on recurrent molecular aberrations from the MET500 study^111^. To map copy number segments to genes, the *GenomicRanges* R package was used to convert segment data into genomic ranges, which were annotated with gene-level data using the *annotatr* R package. A list of 723 known cancer driver genes was sourced from the COSMIC database. Each sample was assigned a G0 score as described in Wiecek et al.^36^ using our previously developed *QuieScore* R package (https://github.com/dkornai/QuieScore). Tumours with a G0 score above 1.5 were classified as slow-cycling, while those with scores below −1.5 were considered fast-cycling. Logistic regression was employed to predict high levels of proliferation versus G0 arrest in TCGA and MET500 cohorts, with model selection performed using the `stepAIC()` function from the *MASS* R package. For model building, samples were split into training (75%) and testing (25%) datasets using the `createDataPartition()` function. A 10-fold cross-validation procedure was implemented using the `trainControl()` function from the *caret* R package. The logistic regression model trained on the MET500 dataset achieved an accuracy of 62.5% with an AUC of 0.66, whilst the model trained on TCGA dataset has an accuracy of 72.7%, with an AUC of 0.78.

### Gene enrichment analysis

Hallmark gene set enrichment analysis on single cells was performed using the *escape* R package. Visualisation of the heatmap was done with the *dittoSeq* R package. Gene Ontology (GO) terms related to DNA replication and the unfolded protein response, as well as the *HALLMARK_P53_PATHWAY* and *HALLMARK_EPITHELIAL_MESENCHYMAL_TRANSITION* gene sets from MSigDB^112^ were assessed by *scDECAF* R package^113^. The top GO terms associated with the gene regulatory network modules were identified using the *enrichR* R package with the *GO_Biological_Process_2023* gene sets. The EMT score was calculated as described in Malagoli-Tagliazucchi et al.^62^. Endoplasmic reticulum stress-related pathways scores were computed using the *irGSEA* R package^114^, with curated gene signature lists as input. Dormancy signatures were taken from Ren et al.^114^, Ruth et al.^115^, Cheng et al.^116^, Kim et al.^117^ and Montagner et al.^43^.

### Cell-cell interactions

#### TME-receiver cell interactions were assessed by two methods

##### (1) Estimation of putative ligands and downstream genes in the receiver population

The estimation of ligands in the TME potentially driving gene expression in G0 arrest or fast-cycling cells involved utilising NicheNet 2.0 R package^69^. Differentially expressed genes with a log2 fold-change greater than 0.5 and an adjusted p-value lesser than 0.05 were considered for ligand prioritisation in the receiver population. Background genes were defined as genes expressed in the receiver population (at least 5% of the cells). Ligands were not constrained to be expressed solely by putative sender cells within the dataset, allowing for the assessment of TME ligands produced by non-immune cells such as tumour cells or the stromal compartment. The prioritisation of ligands was based on the area under the precision-recall (AUPR) scores, indicating whether a gene significantly correlated with their respective ligand-target regulatory potential. The ligand-target regulatory potential reflects the strength of association between a ligand and a target gene based on prior knowledge, presented in NicheNet’s integrated weighted networks. Finally, the predicted interaction network map was plotted using the *DiagrammeR* R package.

##### (2) Estimation of ligand-receptor pairs

To infer ligand-receptor pairs between the TME and receiver cells regardless of their effects on downstream gene expression, the differentially expressed interaction module of CellPhoneDB^75^ was employed, with a threshold of 0.1 percent gene expression in each cell. This method utilises a list of differentially expressed genes sorted by adjusted p-values calculated by Seurat’s `FindAllMarkers()` function for each cell type.

In spatial transcriptomics, we applied the LIANA+ package’s CellPhoneDB module^118^ to identify spatially relevant ligand-receptor pairs.

### Gene regulatory networks

Single-cell regulatory network inference and clustering (pySCENIC)^59^ was employed to infer gene regulatory networks from single cells across G0 arrested, fast-cycling and slow-cycling tumours. The inference of gene regulatory networks was based on *hg38 v10nr_clust databases*, focusing on regions around 10 kb, 100 bp downstream, and 500 bp upstream of transcription start sites (TSS), using score and ranking matrices provided by the SCENIC pipeline.

The top 100 GRNs were grouped by cancer subtype for AUCell scoring, and Spearman’s correlation was applied to identify conserved GRN modules. Binding motifs for selected transcription factors were inferred from the *motifs-v10nr_clust-nr.hgnc-m0.001-o0.0.tbl* database.

### Deconvolution of T cell states

A comprehensive human reference T cell atlas^119^ was used to query the T cell states. For projecting CD8 T cell subtypes onto single cells, *ProjecTILs* R package^120^ was employed.

### Spatial analysis

Breast cancer Visium slides were sourced from Barkley et al.^121^ (slides 0-2), 10X Genomics^122^ (slides 3-5) and Wu et al.^35^ (slides 6-12). These data sets were merged into a unified AnnData python format for analysis. Pre-processing, normalisation and cell type deconvolution were carried out as conducted in previous studies^83^. A total of 32,845 spatially profiled spots were analysed. Spots were retained if they showed at least 100 genes with a minimum count of 1 per cell, had over 250 counts per spot, and less than 20% mitochondrial counts per cell. Cellular deconvolution was performed using cell2location^123^ with a scRNA-seq breast cancer dataset from Wu et al.^35^, comprising 100,064 cells from 26 patients across 21 cell types. These included cancer epithelial cells (basal, cycling, Her2, LumA, LumB), B-cells (naïve and memory), CAF subtypes (myCAF-like and iCAF-like), perivascular-like cells (immature, cycling, differentiated), T-cells (cycling, CD 4, CD 8 T cells), cycling myeloid cells, dendritic cells, endothelial cells (ACKR1, CXCL12, RGS5), LYVE1-expressing lymphatic endothelial cells, luminal progenitors, mature luminal cells, macrophages, monocytes, myoepithelial cells, NK cells, NKT cells, and plasmablasts. Tumour cells in the spatial transcriptomics dataset were identified using the STARCH python package, which infers copy number alterations^124^.

Hotspot analysis was conducted using SpottedPy python package^83^, where the EMT, hypoxia and UPR and angiogenic hallmarks^35^, as well as hybrid E/M and semaphorin signalling signatures were scored using scanpy’s `score_genes()` function. G0 arrest and fast-cycling tumour spots were calculated using the gene sets described in Wiecek et al.^36^.

### Drug-niche interactions

To interrogate drug-niche interactions, we utilised drug2cell^87^ to score candidate drugs and their targets with differential correlations to drug sensitivity by leveraging gene expressions in each cell state, then employed our SpottedPy^83^ method to evaluate the spatial distances between drug niches and cell cycle states.

### The large language model

The G0-LM model is an advanced architecture comprising three specialised networks, adapted from the scBERT framework^125^ to address the specific challenges of G0 arrest in single-cell analysis. Its two-stage training pipeline bridges the gap between large pre-trained datasets and the specialised requirements of G0 arrest data^92^. In the first stage, the model’s embedding space is refined using a Low-Rank Adaptation (LoRA) approach^126^, a Parameter Efficient Fine-Tuning (PEFT) strategy that enables adaptation to scRNA-seq data while reducing overfitting risks.

The G0-LM architecture follows that of EMT-LM^92^ and incorporates a multiplication layer that leverages fine-tuned language models to amplify signals from highly expressed genes. Attention vectors generated by these models act as weighted enhancements of raw gene expression data, selectively highlighting relevant cellular features. To optimise model focus, genes with low expression or peripheral relevance to the EMT process are systematically filtered, concentrating learning on critical pathways. For classification, a Multilayer Perceptron (MLP) serves as the final module to yield predictions. Ultimately, this fusion network provides a computational model of cell states under G0 arrest, distilling complex gene expression profiles into accurate cell state outputs.

### Statistical analysis

Groups were compared using a two-sided Student’s *t*-test, Wilcoxon rank-sum test, ANOVA or Kruskal-Wallis test, as appropriate. Adjustments for multiple testing were made using the Benjamini-Hochberg method when necessary. False discovery rates (FDR) were calculated by taking the −log10 of the adjusted *p*-values. Graphs were generated using either the *ggplot2* and *ggpubr* R packages or the *matplotlib* and *seaborn* python libraries.

## Data availability

The single-cell RNA sequencing datasets used in this study are publicly available through the Curated Cancer Cell Atlas of the Weizmann Institute of Science (https://www.weizmann.ac.il/sites/3CA/breast), with raw files accessible via the Gene Expression Omnibus (GEO) under accession numbers GSE148673^38^ and GSE161529^39^, and through the ArrayExpress database of EMBL-EBI under accession number E-MTAB-8107^37^. Pre-processed and annotated single-cell Seurat objects are available on Zenodo (DOI: 10.5281/zenodo.14001194). Gene sets for calculating G0 arrest cell cycle score are available at https://github.com/secrierlab/CancerG0Arrest. The human CD8 T cell reference dataset is provided in R dataset format and can be downloaded from Figshare (DOI: 10.6084/m9.figshare.41414556). Processed spatial transcriptomics data is also publicly available on Zenodo (DOI: 10.5281/zenodo.10371890).

## Code availability

All scripts used in this study are publicly available at https://github.com/secrierlab/G0-breast-cancer-atlas. The G0-LM model is available at https://github.com/secrierlab/G0-LM.

## Acknowledgements

M.S., C.C. and S.P. were supported by a UKRI Future Leaders Fellowship (MR/T042184/1). E.W. was supported by a studentship award from the Health Data Research UK-The Alan Turing Institute Wellcome PhD Programme in Health Data Science (218529/Z/19/Z). Work in M.S.’s lab was supported by a BBSRC equipment grant (BB/R01356X/1) and a Wellcome Institutional Strategic Support Fund (204841/Z/16/Z). J.L. was supported by BBSRC grants BB/T013273/1 and BB/W014890/1.

We would like to acknowledge Dr Andrew Holding for his input on visualising hotspot overlaps in the spatial transcriptomics slides, and Dr Kevin Litchfield and Dr Andrea Castro for their help investigating the therapeutic relevance of the G0 arrest signature.

## Authors’ contributions

M.S. designed and supervised the study. C.C. and M.S. designed the experiments and interpreted the data. C.C. performed all analyses, except for the following: E.W. developed SpottedPy, processed the spatial data and conducted the initial SpottedPy analyses; T.C. performed logistic regression analyses of genomic alterations in fast/slow proliferating tumours in bulk datasets from TCGA and MET500, as well as enrichment analyses of genomic changes in G0 arrest and fast-cycling single cell data; S.P. developed the G0-LLM model and analysed the corresponding data. J.L. provided a curated signature list for UPR and led the interpretation of the proteostasis-linked analyses. C.C., M.S., S.P. and J.L. wrote the manuscript. All authors read and approved the manuscript.

## Ethics declarations

All datasets employed in this study are publicly available and comply with ethical regulations, with approval and informed consent for collection and sharing already obtained by the relevant organizations.

