## Supplementary Material for "Balancing tumour proliferation and sustained cell cycle arrest through proteostasis remodelling drives immune niche compartmentalisation in breast cancer"

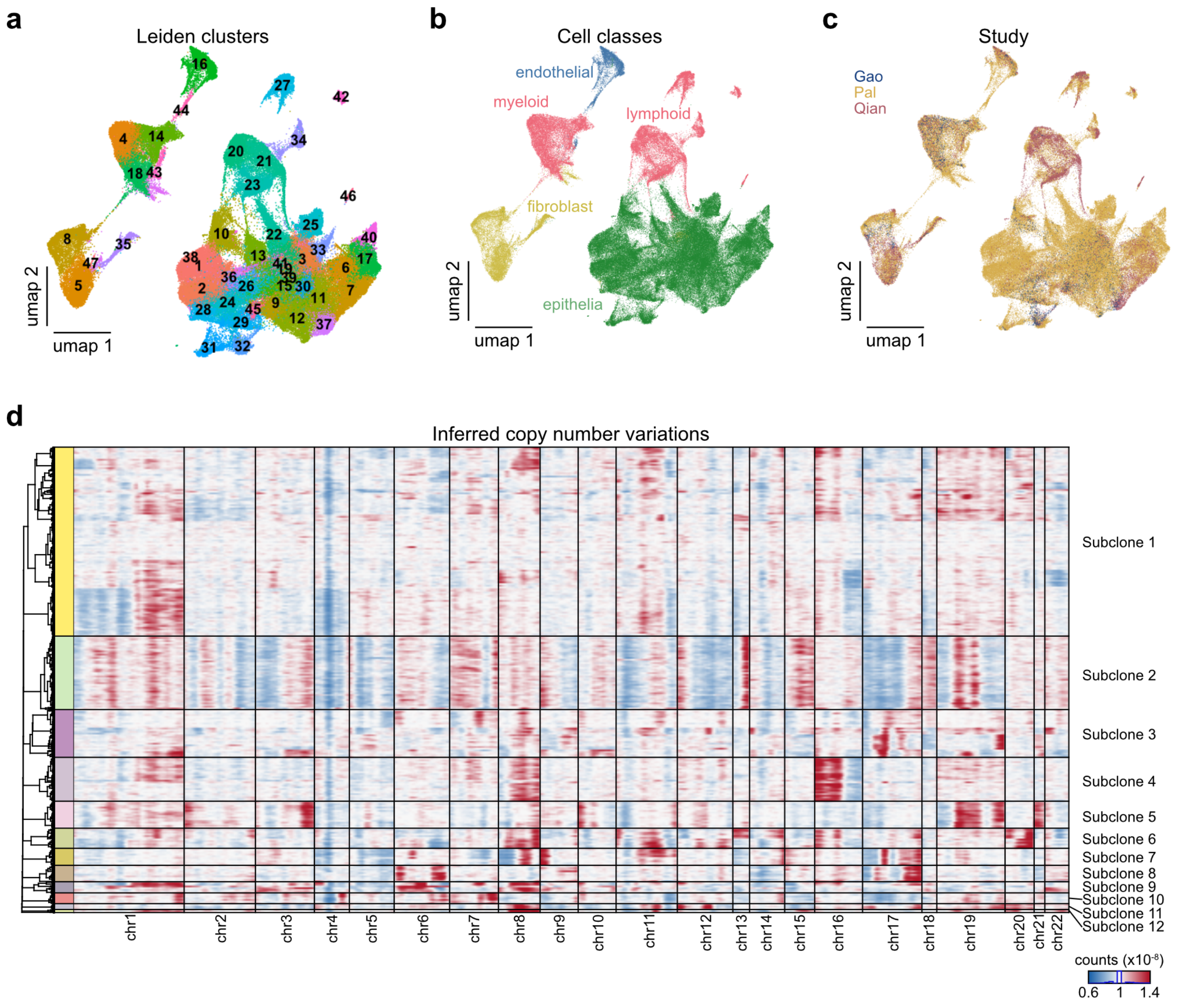


**Extended Data Fig. 1 | Characteristics of the integrated dataset.**

**a-c** UMAP plots show the distribution of Leiden clusters **(a)**, cell classes **(b)**, and study sources **(c)** within the integrated dataset. **(d)** Inferred copy number alterations in epithelial cells.

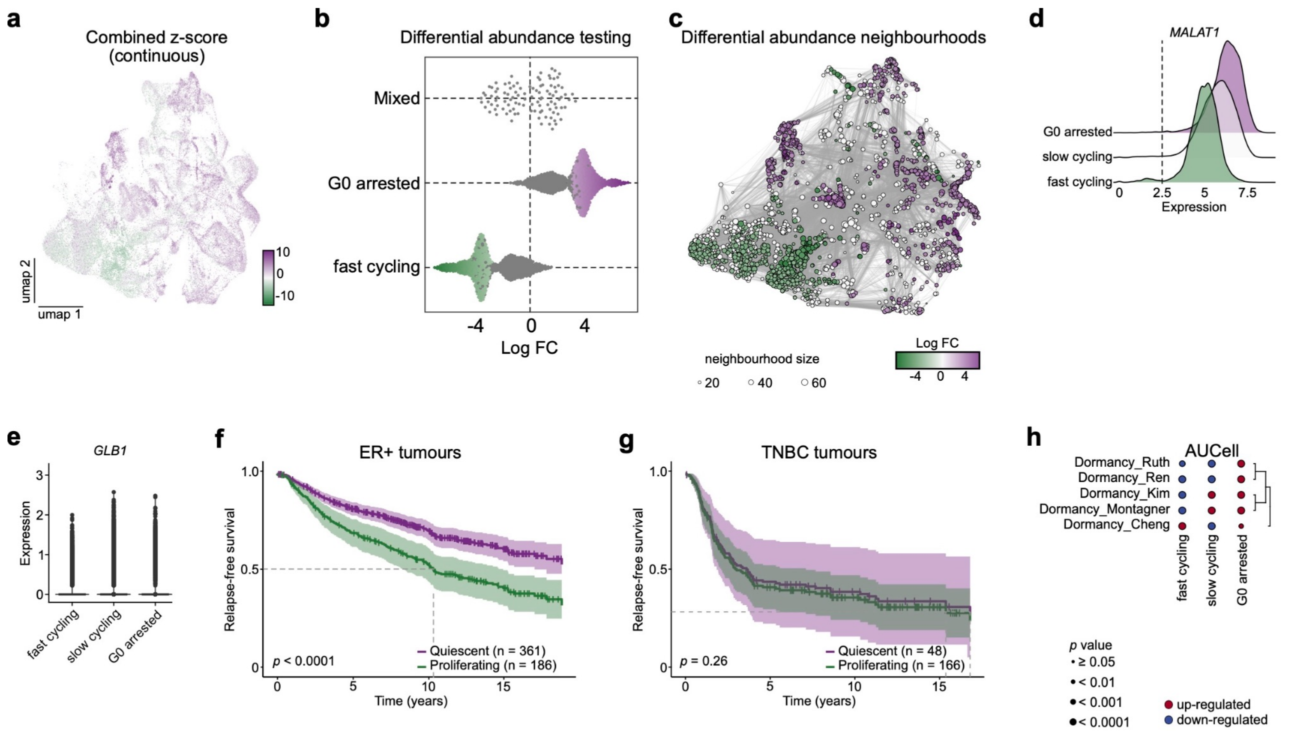


**Extended Data Fig. 2 | Cell cycle states. a** Continuous G0 arrest score in malignant cells. **b-c** Differential abundance test **(b)** and neighbourhood map **(c)** highlighting distinct cell cycle states. **d-e** Expression of *MALAT1* **(d)** and *GLB1* **(e)**. **f-g** Kaplan-Meier survival plots for ER+ **(f)** and TNBC tumours **(g)**. **h** AUCell scores for different dormancy signatures comparing cell cycle states.

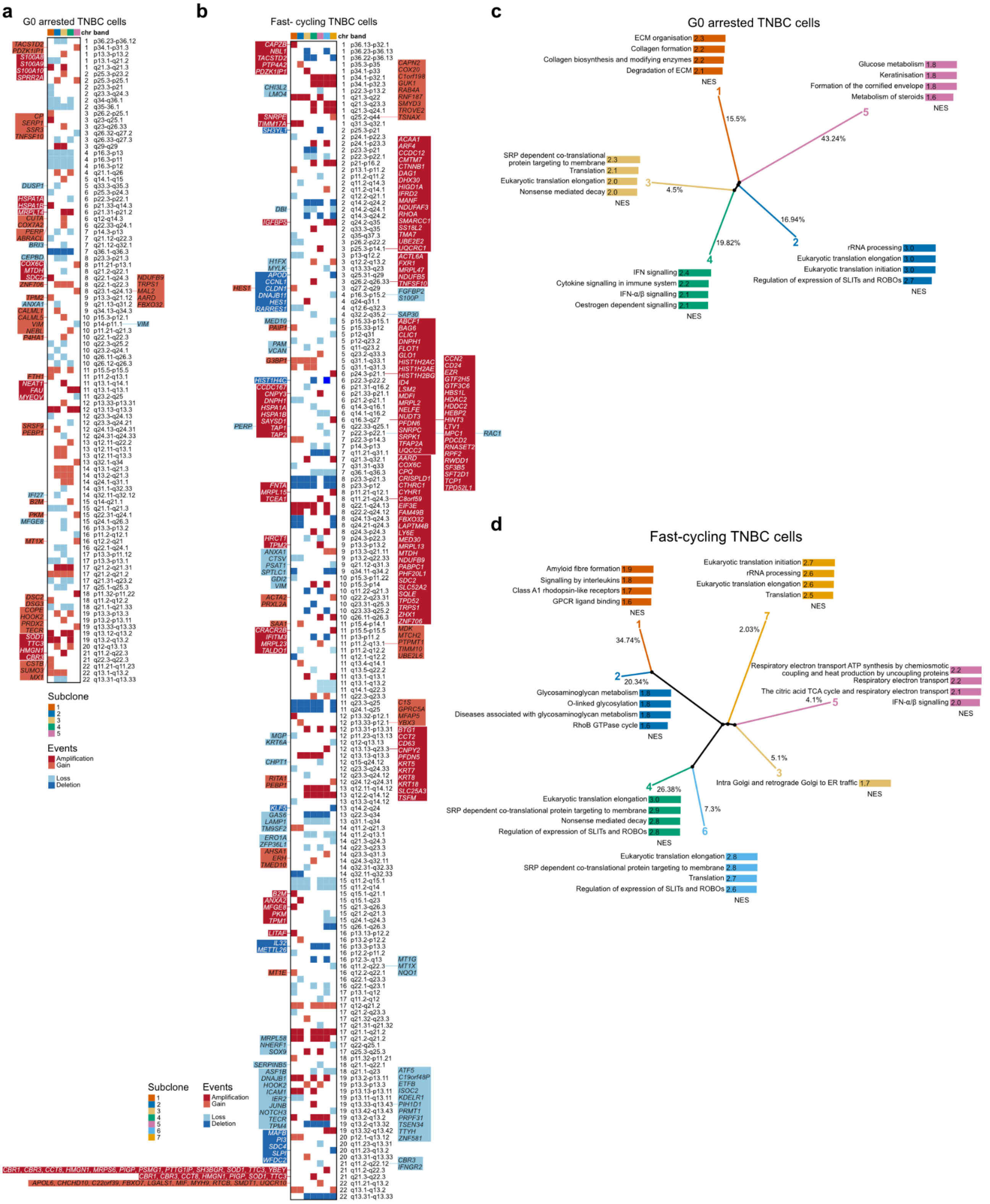


**Extended Data Fig. 3 | Genomic differences between G0 arrested and fast-proliferating tumours. a** Chromosomal arm-level alterations in G0 arrested TNBC subclones, highlighting shared and unique genomic changes between subclones, as inferred by SCEVAN. Differentially expressed genes between G0 arrested and fast-cycling cells at each chromosomal location are highlighted on the side. **b** Chromosomal arm-level alterations in fast-proliferating TNBC subclones, illustrating key genomic differences compared to G0 arrested states. **c** Clone tree of G0 arrested TNBC tumour cells with subclones colour-coded by Reactome pathway enrichment of the correspondingly altered genes. **d** Clone tree of fast-proliferating TNBC tumour cells, colour-coded similarly by Reactome pathway enrichment.

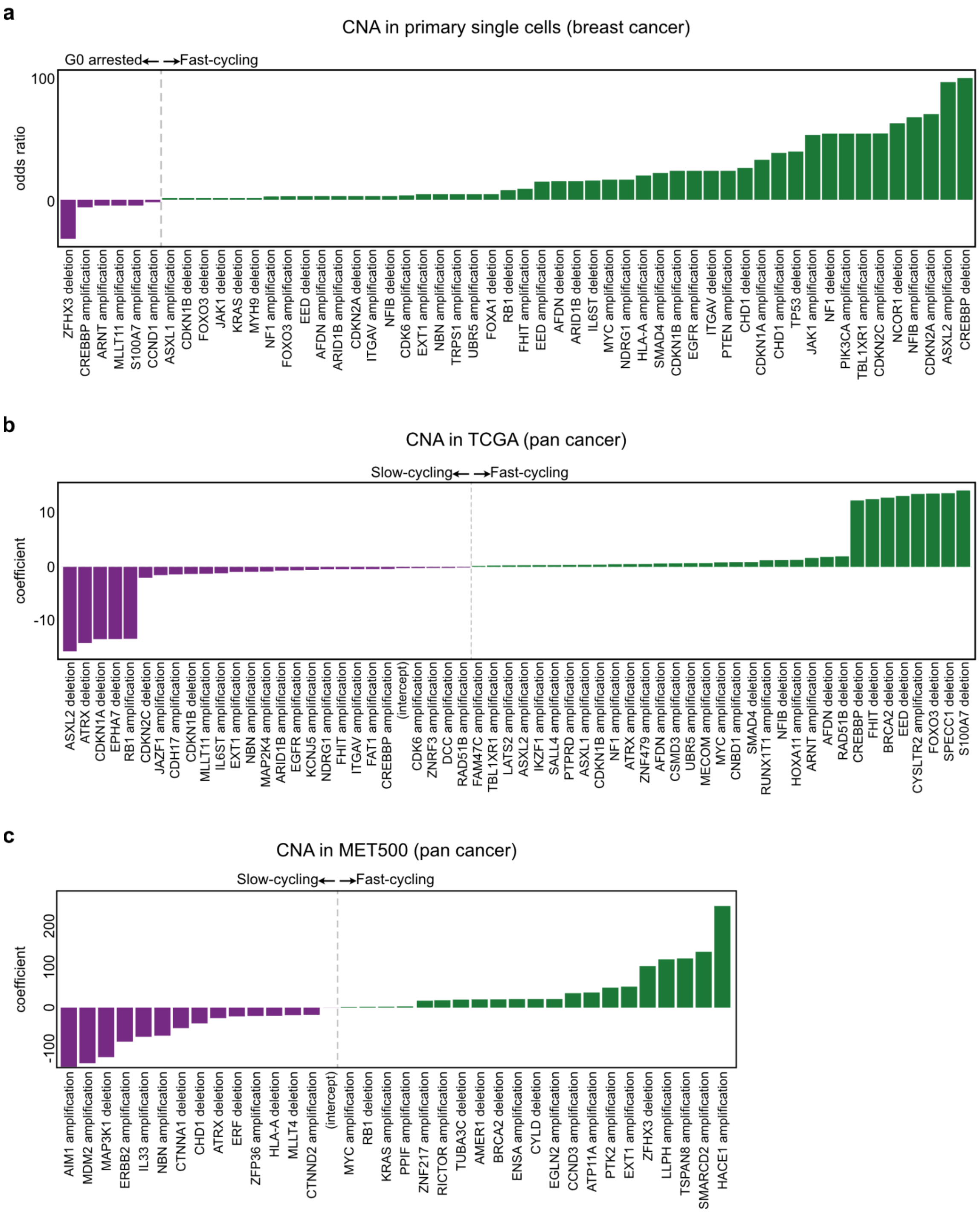


**Extended Data Fig. 4 | Genomic alterations in cancer drivers linked with tumour cell cycle state. a** Copy number changes in cancer drivers differentially present in G0 arrested (purple) or fast cycling (green) cells in primary single cell breast cancer cohorts, calculated using Fisher's exact tests. **b-c** Genomic changes in cancer drivers linked with slow cycling versus fast cycling TCGA primary tumours **(b)** and MET500 metastatic tumours **(c),** inferred based on logistic regression models.

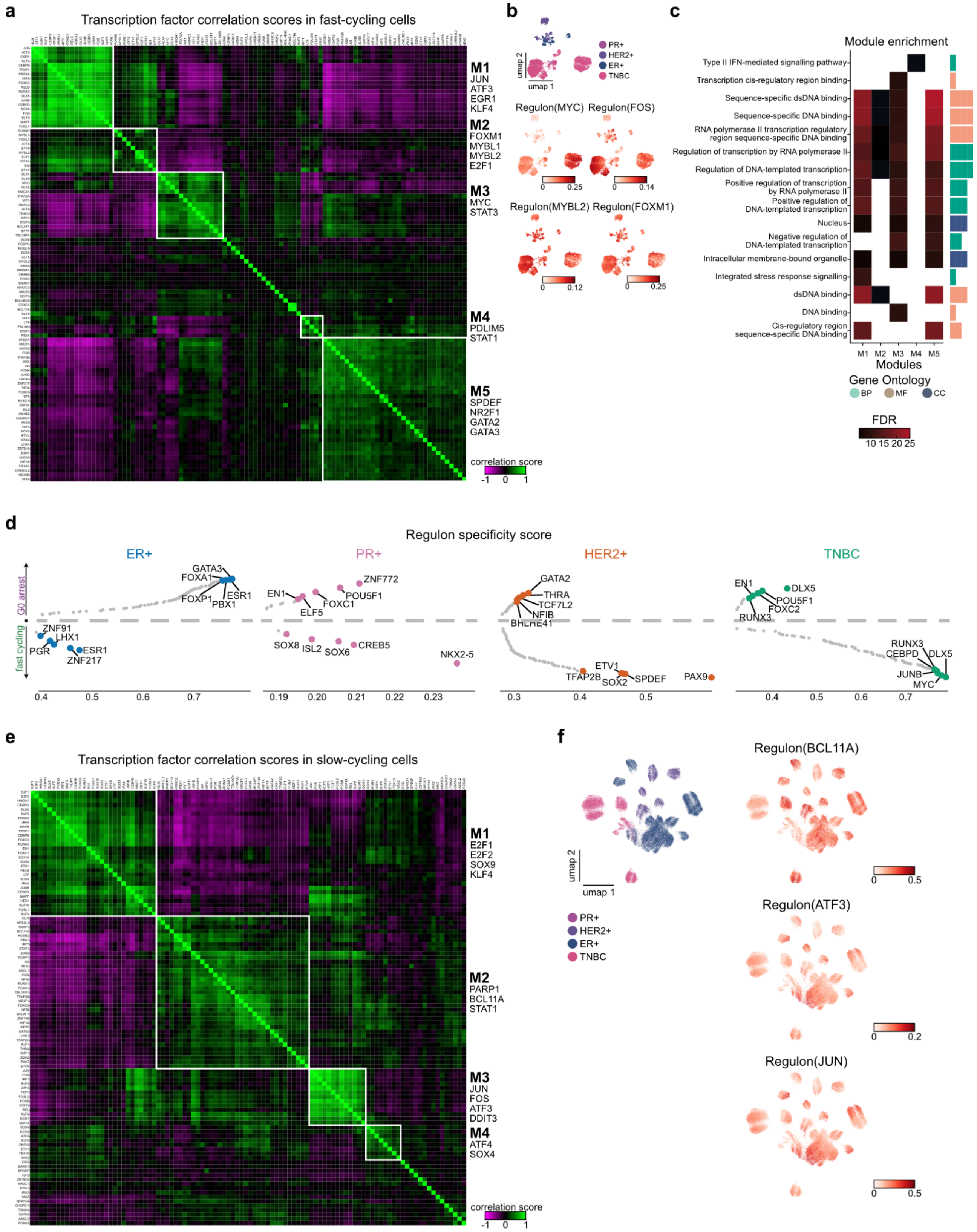


**Extended Data Fig. 5 | Gene regulatory networks modulating G0 arrest in fast-cycling and slow-cycling cells. a** Heat map of correlation scores for the top 100 transcription factors (TFs) in fast-cycling cells, with modules (M1–M5) highlighted. **b** UMAP plot of fast-cycling cells showing regulon activities of selected TFs. **c** Pathway enrichment for every module discovered in fast-cycling cells. **d** Regulon specificity scores (x axis), comparing the top TFs between G0 arrested and fast-cycling (y axis) cells for each breast cancer subtype. **e** Heat map of correlation scores for the top 100 TFs in slow-cycling cells, with modules (M1–M4) highlighted. **f** UMAP plot of slow-cycling cells with regulon activities of selected TFs.

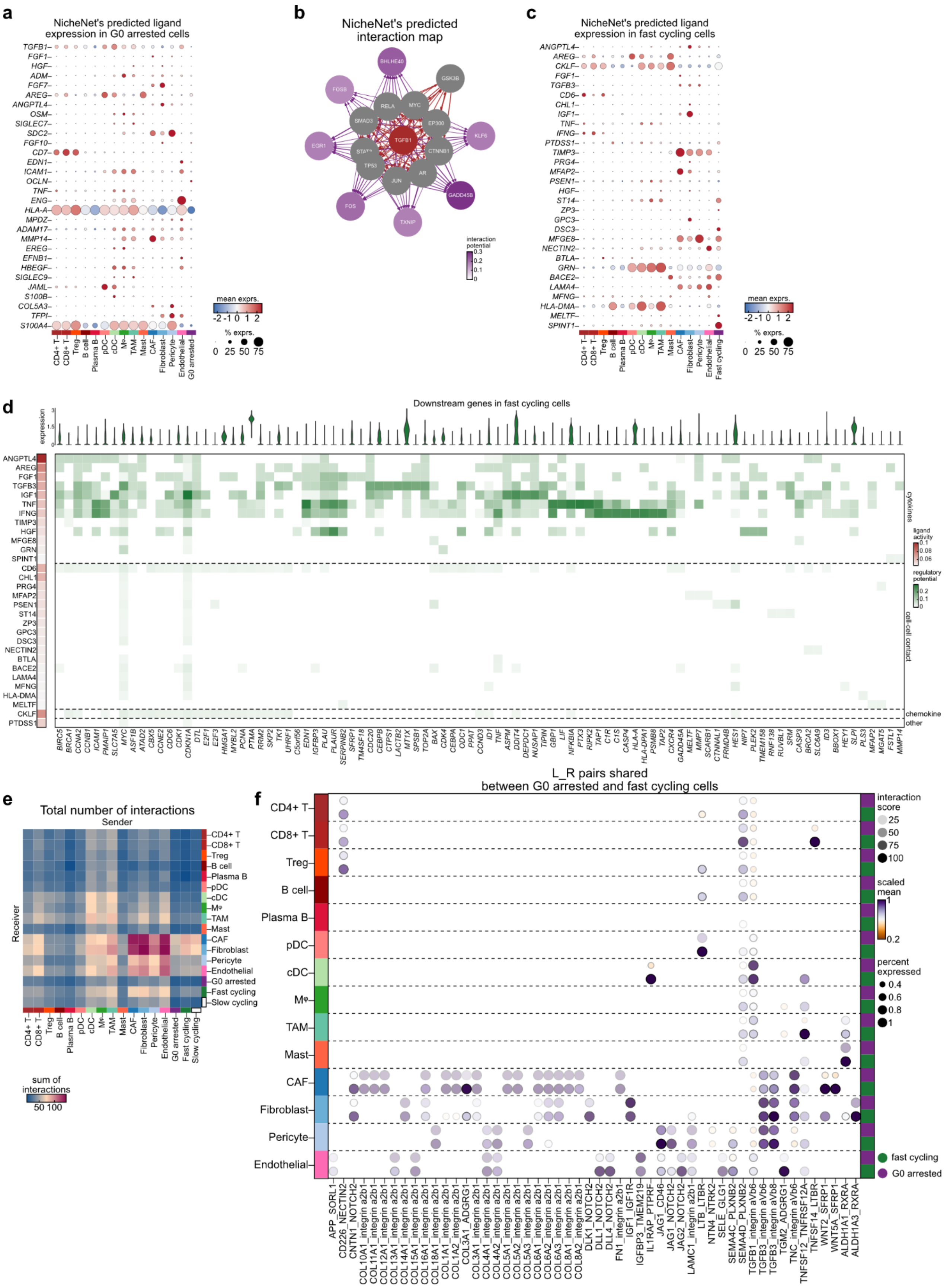


**Extended Data Fig. 6 | Cell-cell interactions in the TME. a** Bubble plot of predicted ligand expression in the TME affecting G0 arrested cells. Rows correspond to ligands expressed by G0

arrested cells, columns indicate the receiving cells in the TME. **b** NicheNet's predicted ligand-target gene expression interaction map in G0 arrested cells, where colours in target genes denote interaction potential with TGFB1. **c** Bubble plot of predicted ligand expression in the TME affecting fast-cycling cells. Rows correspond to ligands expressed by fast cycling cells, columns indicate the receiving cells in the TME. **d** The top 30 ligands expressed by cells in the TME and their target genes in fast-cycling cells, as inferred by NicheNet. Rows highlight ligands on immune and stromal cells, alongside their activity depicted by a white-red colour gradient. The ligands are grouped manually into biologically relevant categories, annotated to the right. Columns depict downstream targets in fast cycling cancer cells. Violin plots (top) show target gene expression in fast-cycling cells. **e** Total number of ligand-receptor interactions in the TME. **f** Shared ligand-receptor pairs between fast-cycling and G0 arrested cells within the TME. The rows depict the sender cells in the TME, the columns highlight shared ligand-receptor pairs with the receiver tumour cell, as inferred by CellPhoneDB.

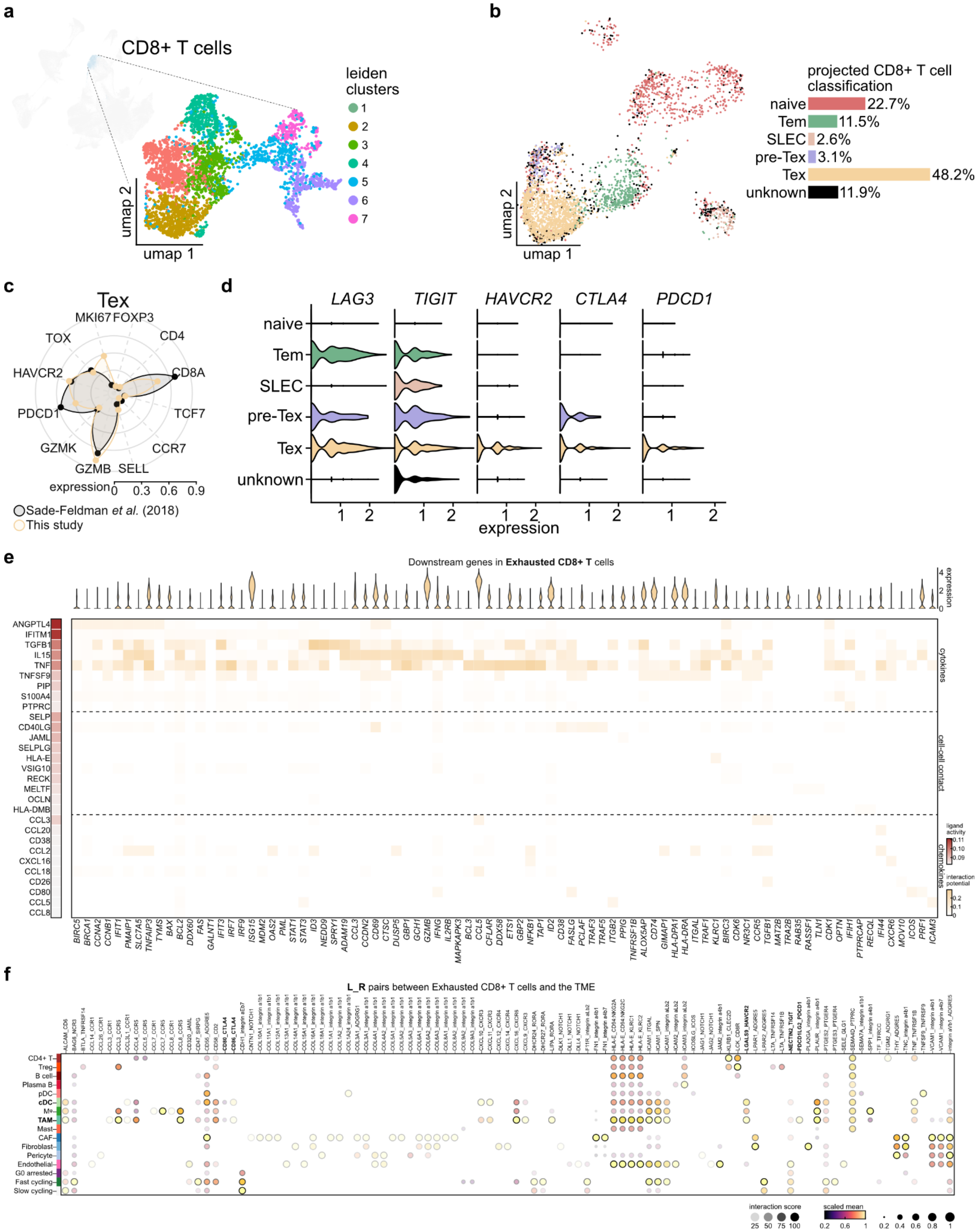


**Extended Data Fig. 7 | CD8 T cell subtype characterisation. a** UMAP of CD8 T cell subsets identifying seven Leiden clusters. **b** CD8 T cell label transfer from a reference T cell atlas. **c** Spiderweb plot comparing exhausted T cell marker expression with the reference dataset. **d** Violin plots of exhausted T cell marker expression across CD8 T cell subtypes. **e** Heat map showing the interaction potential of the 30 top prioritised ligands in fast-cycling cells (rows) and target genes in exhausted T cells (columns). The predicted ligand activity by biological categories is shown to the left. Violin plots (top) show expression levels of target genes in exhausted T cells. **f** Ligand-receptor pairs between exhausted T cells (rows) and the TME (columns).

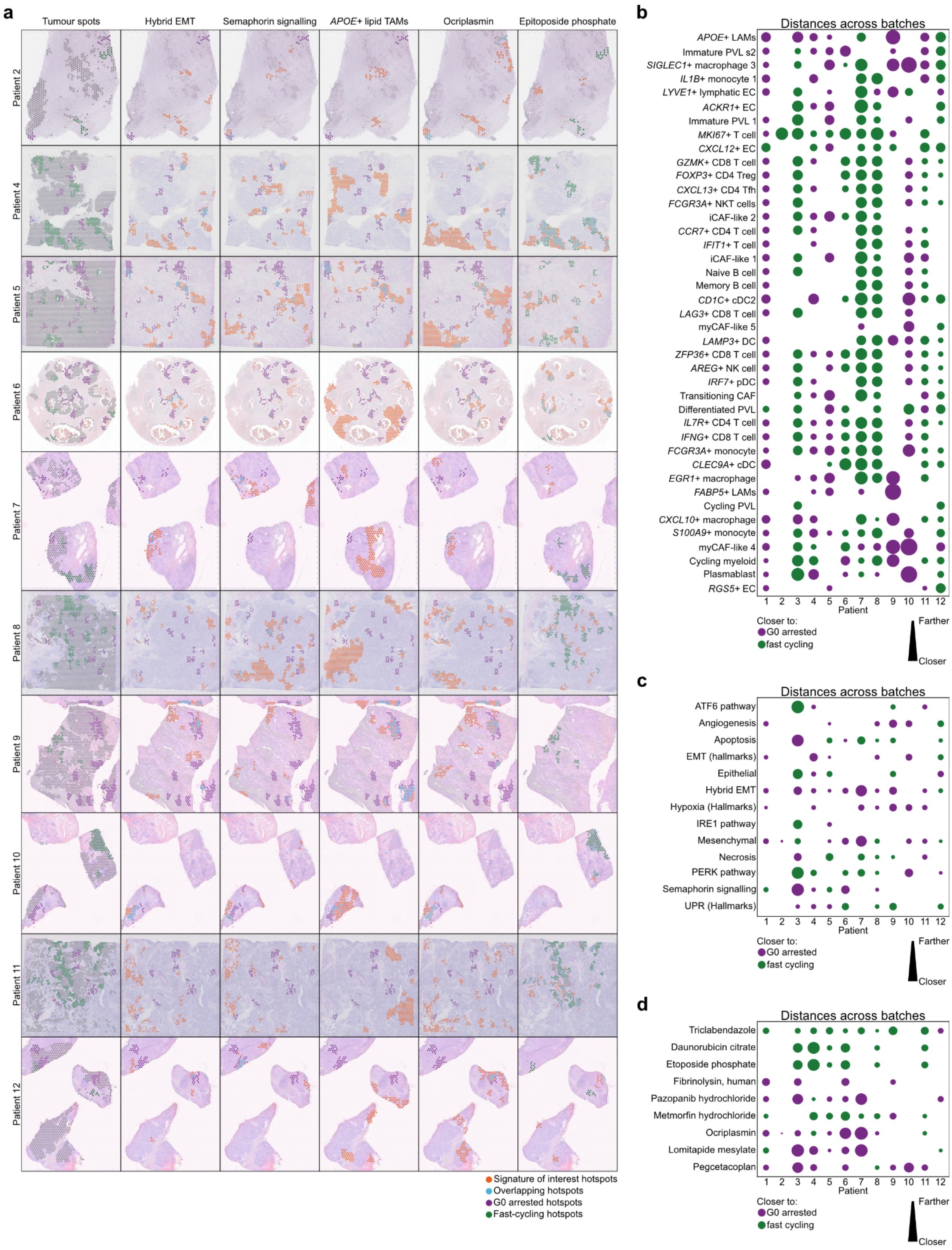


**Extended Data Fig. 8 | Spatial organisation of cancer hotspots in breast primary tumours.**
**a** Spatial transcriptomics images displaying hotspots identified by SpottedPy in ten patient samples. Hotspots are colour-coded by cell cycle state: green (fast-cycling), grey (slow-cycling), and purple (G0 arrested). Orange hotspots highlight the signature or cell type of interest, as specified in each panel. Teal hotspots indicate overlapping regions between G0 arrested/fast-cycling hotspots and the

signature/cell type/drug interaction of interest. **b-d** Bubble plots showing relative median Euclidean distances between defined hotspots, illustrating the proximity of cell type-specific hotspots **(b),** signature-specific hotspots **(c)** and drug interaction hotspots **(d)** to either G0 arrested or fast-cycling hotspots.

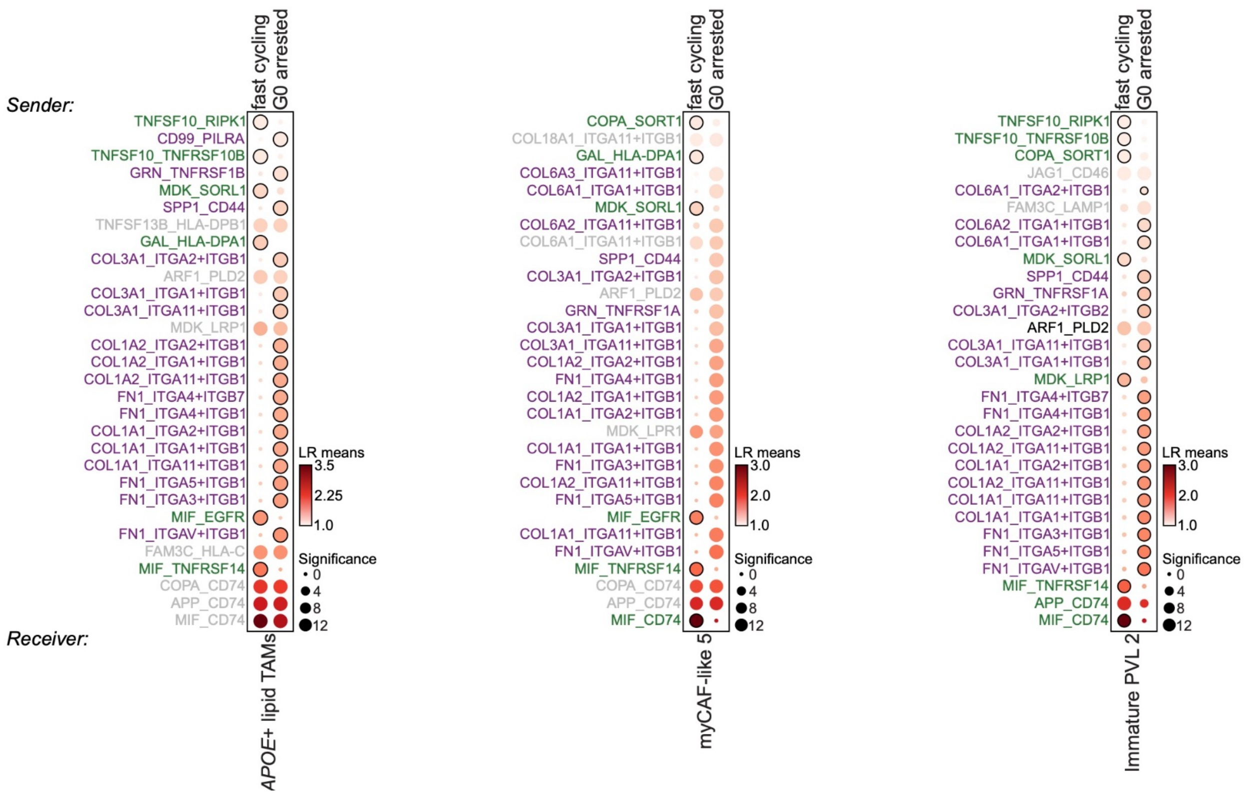


**Extended Data Fig. 9 | Ligand-receptor analysis.** Ligand-receptor interaction analysis between tumour cells as senders and *APOE*+ lipid-associated TAMs, myCAF-like and immature PVL cells as receivers, respectively, in relation to G0 arrested and fast-cycling hotspots. Purple indicates ligand-receptor interactions unique to G0 arrested tumour cells, green indicates those unique to fast cycling cells.

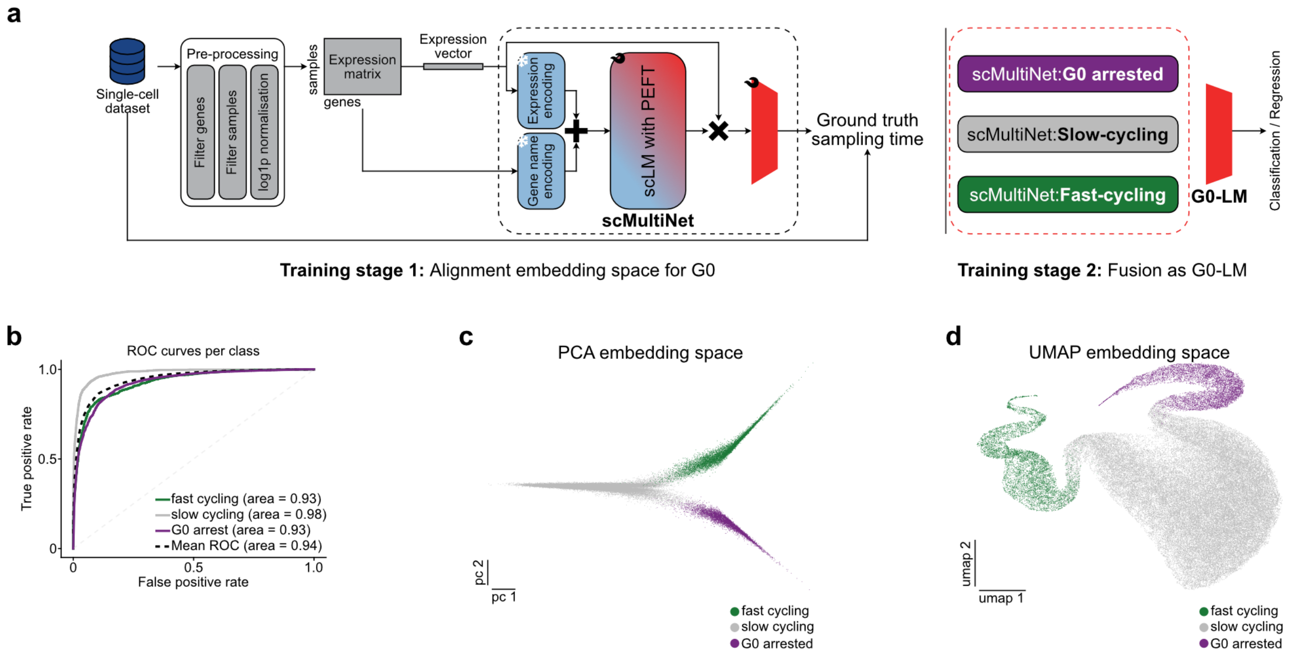


**Extended Data Fig. 10 | G0-LM classification pipeline and performance. a** G0 arrest and proliferation state classification pipeline in scRNA-seq data. The first training phase involves binary classification training for each state separately using the Parameter-Efficient Fine-Tuning (PEFT) method. This allows the original pre-trained model's feature space to better represent G0 arrest/proliferation. The final G0-LM model is obtained by fusing the three binary classification models for the individual states (G0 arrest, slow cycling and fast cycling). **b** Receiver operating characteristic (ROC) curves per cell cycle category displaying performances of G0-LM. **c-d** PCA **(c)** and UMAP **(d)** embedding spaces of the cell cycle categories.
